# Hemispheric Asymmetries of Individual Differences in Functional Connectivity

**DOI:** 10.1101/2022.04.08.487658

**Authors:** Diana C. Perez, Ally Dworetsky, Rodrigo M. Braga, Mark Beeman, Caterina Gratton

## Abstract

Resting-state fMRI studies have revealed that individuals exhibit stable, functionally meaningful divergences in large-scale network organization. The locations with strongest deviations (called network ‘variants’) have a characteristic spatial distribution, with qualitative evidence from prior reports suggesting that this distribution differs across hemispheres. Hemispheric asymmetries can inform us on constraints guiding the development of these idiosyncratic regions. Here, we used data from the Human Connectome Project to systematically investigate hemispheric differences in network variants. Variants were significantly larger in the right hemisphere, particularly along the frontal operculum and medial frontal cortex. Variants in the left hemisphere appeared most commonly around the temporoparietal junction. We investigated how variant asymmetries vary by functional network and how they compare with typical network distributions. For some networks, variants seemingly increase group-average network asymmetries (e.g., the group-average language network is slightly bigger in the left hemisphere and variants also appeared more frequently in that hemisphere). For other networks, variants counter the group-average network asymmetries (e.g., the default mode network is slightly bigger in the left hemisphere, but variants were more frequent in the right hemisphere). Intriguingly, left- and right-handers differed in their network variant asymmetries for the cinguloopercular and frontoparietal networks, suggesting that variant asymmetries are connected to lateralized traits. These findings demonstrate that idiosyncratic aspects of brain organization differ systematically across the hemispheres. We discuss how these asymmetries in brain organization may inform us on developmental constraints of network variants, and how they may relate to functions differentially linked to the two hemispheres.

## 1. Introduction

Resting-state functional Magnetic Resonance Imaging (***rs-fMRI***) has become a powerful tool for studying the underlying functional architecture of the brain by examining correlated intrinsic activity between brain regions ((Biswal et al., 1995); for a review see (Buckner et al., 2013; Power et al., 2015)). This approach has resulted in a robust description of functional networks that are linked to cognitive and sensorimotor processes (Doucet et al., 2011; Power et al., 2011; Yeo et al., 2011). The majority of research on this topic has used the traditional approach of grouping data together from many participants. However, there is individual variability in functional network organization, particularly in association cortex (Mueller et al., 2013). Thus, unique features at the individual level are blurred as a result of averaging across brains with varying network topography.

Recent efforts to map network architectures at the individual level have used ***precision fMRI***, an approach in which experimenters collect extended amounts of rs-fMRI data across multiple sessions for each subject to obtain highly reliable individualized measurements (Braga & Buckner, 2017; Gordon, Laumann, Gilmore, et al., 2017; Greene et al., 2020; Laumann et al., 2015). While precision fMRI studies suggest that functional networks follow roughly the same organizational principles across participants, each person also exhibits idiosyncratic features: localized regions where an individual’s functional connectivity pattern differs markedly from the group average (i.e. the region shows a pattern of connectivity that is more correlated with a different network than what would typically be found at that location) (Gordon, Laumann, Gilmore, et al., 2017; Laumann et al., 2015; Seitzman et al., 2019). We refer to these regions of individual difference as ***network variants*** (henceforth also referred to as variants). Network variants can arise for two reasons: either due to relatively local shifts in the borders between networks (‘border variants’) or due to more dramatic changes in network organization far from the typically associated network (‘ectopic intrusions’; (Dworetsky et al., 2021). While variants (of both types) are widespread, individuals differ in the precise location, size, network characteristics and forms of network variants they exhibit (Dworetsky, Seitzman, Adeyemo, Smith, et al., 2021; Seitzman et al., 2019). Initial studies demonstrate that individual differences in brain network organization relate to individual differences in cognition measured outside of the scanner (Bijsterbosch et al., 2018; R. Kong et al., 2019; Seitzman et al., 2019) as well as differences in task evoked responses during MRI (Seitzman et al., 2019).

Network variants have been shown to be stable over time and to exhibit a characteristic spatial distribution—they are most commonly observed in frontal and temporo-parietal cortex in regions overlapping higher-order cognitive networks in the group average (Seitzman et al., 2019). Additionally, network variants are often associated with higher-level attention, default, and language networks, and rarely associated with networks involved in sensorimotor processes. Intriguingly, by visual examination, these individual differences appear to be relatively lateralized, with seemingly more variants appearing in the right hemisphere (see Fig. 3A in (Seitzman et al., 2019)). Factors including evolutionary (Buckner & Krienen, 2013), genetic (Anderson et al., 2021), and experiential variables (Newbold et al., 2020) have been proposed to guide the organization of closely-juxtaposed networks as they fractionate and specialize in the cortex (DiNicola & Buckner, 2021). However, the constraints that influence how and where individual differences in these networks occur are currently largely unknown.

Examining hemispheric asymmetries in network variants may inform how individual differences in functional brain organization arise. While there is a large degree of homology between the two hemispheres, several asymmetries have been reported in the human brain, ranging from functional to anatomical and cytoarchitectonic in nature (see (Toga & Thompson, 2003) for a review). Macrostructural asymmetries are observed in a number of locations, including the trajectory of the Sylvian fissure (Hou et al., 2019), the volume of the temporal plane (Geschwind & Levitsky, 1968; Steinmetz, 1996), and in frontal and occipital petalia (X.-Z. Kong et al., 2018, 2019). In addition, the two hemispheres have been differentially tied to specific functions. Perhaps the most notable examples for asymmetric function are language and visuospatial attention. While the right hemisphere makes important contributions to language processing (Jung-Beeman, 2005; Lindell, 2006), most individuals exhibit left-hemispheric dominance for core language function (Breier et al., 2000; Stippich et al., 2003), especially language production (Lidzba et al., 2011). In contrast, the majority of the population exhibits right-hemisphere bias for visuospatial processing (Cai et al., 2013; Weintraub & Mesulam, 1987). Asymmetries can also be present in behavioral traits like handedness, auditory perception, and motor preferences (Toga & Thompson, 2003). Functional brain networks also show hemispheric differences in group-averages (Gotts et al., 2013; Wang et al., 2014). This is especially true in the default mode network, which shows a left-hemispheric specialization (strong within-hemispheric interactions), the ventral and dorsal attention networks which show right-hemispheric specialization, and the frontoparietal network, which shows specialization in the two hemispheres, but couples with different networks in each hemisphere (Wang et al., 2014). The language network is often hard to detect in group-averages due to strong variation across people (Braga et al., 2020; Fedorenko, 2012), but individual data suggest that it is relatively left-lateralized.

These previous investigations show that hemispheric asymmetries exist at every level of brain organization. Here, we ask how individual differences in network organization vary across the two hemispheres by studying asymmetries in the properties of network variants. Such asymmetries can help our understanding of how individual differences arise and may elucidate important aspects of the relationship between brain organization and cognitive function.

In this project, we first examine hemisphere-wide asymmetries in the extent and size of idiosyncratic variant regions in a large group of young adult subjects from the Human Connectome Project. These asymmetries in the general features of variants would suggest differences in the overall developmental constraints of the two hemispheres. For example, a rightward lateralization in the frequency of variants would indicate a more variable architecture of the right hemisphere compared to the left hemisphere across individuals. Speculatively, this may indicate that the right hemisphere is less constrained by developmental programs and more malleable to experience-dependent re-shaping. Next, we focus on specific locations and networks exhibiting hemispheric asymmetries and study how these asymmetries in network variants relate to group-average network asymmetries. Lastly, we investigate a potential link between individual differences in functional connectivity and behavior by looking at the relationship between network variant asymmetries and handedness.

## 2. Materials and Methods

### 2.1 Datasets and overview

Two independent and publicly available datasets were used for these analyses: The Human Connectome Project (Van Essen, Ugurbil, et al., 2012) and the Midnight Scan Club (Gordon, Laumann, Gilmore, et al., 2017). These datasets were used due to their relatively high amounts of low-motion data per subject, which allows individual-specific measures of functional networks to reach high reliability levels. Specifically, network variant measurements of the cortex can reach high reliability (test-retest r > 0.8) with about >45 minutes of good quality data or more (Kraus et al., 2021; Laumann et al., 2015).

Our primary analyses were based on a subsample (N = 384) of unrelated subjects with > 45 min. of low-motion data from the HCP dataset. The HCP 1200-subject release includes one hour of resting-state data collected across two sessions for 1018 subjects (female = 546) between the ages of 22-35, including data from twins and non-twin siblings (for more details on the demographic breakdown of this sample, the reader is referred to Van Essen et al., 2013). Exclusion criteria for the HCP 1200-subject release included removing subjects who did not complete the study, and subjects who did not have at least 45 minutes of high-quality resting-state data after motion censoring (single subject measurements of network variants achieve good reliability with more than 45 min. of low motion data; (Kraus et al., 2021)). This resulted in a subsample of 752 participants (female = 423; see Seitzman et al., 2019 for more details). If more than one member of a given set of siblings met the previous criteria, the subject with the most data was selected and their sibling(s) were excluded from most analyses, resulting in the subsample of 384 unrelated subjects (female = 210). Note that, in order to increase the number of left-handed individuals, for the left- vs. right-hander comparisons, we used the full subsample of 752 subjects from the HCP 1200 release, including related individuals. For the handedness group comparisons, subjects were categorized into three groups based on their Edinburgh Handedness Inventory (EHI) scores (Oldfield, 1971), resulting in 40 left handers (scores between -100 and - 41), 670 right handers (scores between 41 and 100), and 42 ambidexters (scores between -40 and 40).

The MSC was used as a replication dataset to confirm the findings of the HCP in a highly sampled precision-fMRI dataset. The MSC includes five hours of resting-state BOLD data from 10 unrelated subjects collected across ten sessions. Due to high motion and drowsiness, one subject was excluded from analyses (Gordon, Laumann, Gilmore, et al., 2017; Laumann et al., 2015). Thus, data from 9 subjects (female = 4, ages 24-34, all right handers) were used to replicate comparisons of spatial distribution of variants.

An additional dataset of 120 neurotypical young adults, the WashU 120 dataset (female = 60, mean age 24.7 ± 2.4 years; (Power et al., 2013)), was used as the group average from which canonical network templates were derived and to which individual-specific connectivity maps were compared to define network variants.

### 2.2 MRI acquisition parameters

The HCP dataset was collected on a Siemens 3T Skyra with a 32-channel head coil. T1-weighted (256 sagittal slices, TR = 2400 ms, TE = 2.14ms, 3D MPRAGE sequence) and T2-weighted (256 sagittal slices, TR =3200ms, TE = 565ms, Siemens SPACE sequence) images were collected for each subject using isotropic 0.7mm voxels (Glasser et al., 2013). High-resolution eyes-open resting-state fMRI data were collected using 2mm isotropic voxels (72 sagittal slices, TR = 720ms, TE = 33ms, multi-band accelerated pulse sequence with multiband factor = 8) (Glasser et al., 2013, 2016). See (Glasser et al., 2013, 2016) for additional detailed information on acquisition in the HCP dataset.

The MSC dataset was collected on a Siemens 3T Trio with a 12-channel head coil. The dataset includes several types of structural data, including four high-resolution T1w images (0.8mm isotropic voxels, 224 sagittal slices, TE = 3.74ms, TR = 2400ms) and four high-resolution T2w images (0.8mm isotropic voxels, 224 sagittal slices, TE = 479ms, TR = 3200ms). Eyes-open resting-state fMRI data were collected using a gradient-echo EPI sequence with 4mm isotropic voxels (36 slices, TE = 27ms, TR = 2200ms) (Gordon, Laumann, Gilmore, et al., 2017).

The WashU 120 dataset was collected on a Siemens 3T Trio with a 12-channel coil and includes a high-resolution T1-weighted image (176 slices, isotropic 1mm voxels, TE = 3.06ms, TR = 2400ms) and eyes-open resting-state BOLD data (TR = 2500ms, TE = 27ms, gradient-echo EPI sequence, 4mm isotropic voxels) (Power et al., 2013).

### 2.3 Preprocessing and functional connectivity processing of functional data

Data were preprocessed identically as in (Seitzman et al., 2019). Briefly, HCP volumetric resting-state time series from each participant were preprocessed as recommended by the minimal preprocessing pipelines (Glasser et al., 2013). Then, the data underwent field map distortion correction, mode-1000 normalization, motion correction via a rigid body transformation within each run, and affine registration of BOLD images to a T1-weighted image and affine alignment into MNI space. The MSC dataset was preprocessed similarly with the following exceptions: this data also underwent slice timing correction and affine alignment was mapped onto Talaraich rather than MNI space.

Resting-state data were further denoised using regression of white matter, cerebrospinal fluid, the global signal, six rigid-body parameters and their derivatives and expansion terms (Friston et al., 1996). High-motion frames were identified using frame-by-frame displacement and censored to remove bias in functional connectivity (Power et al., 2012). Frames with FD > 0.2 for MSC data (Gordon, Laumann, Gilmore, et al., 2017) and filtered FD > 0.1 for the HCP data (Fair et al., 2020) were flagged as being contaminated by motion and removed from analyses. Note that for two MSC subjects (MSC03 and MSC10) the filtered FD (motion parameters low-pass filtered < 0.1 Hz) measure was also used to identify to-be-censored frames in order to address respiratory related artifacts in their FD parameter (Gordon, Laumann, Gilmore, et al., 2017; Gratton et al., 2020).

For both datasets, the first 5 frames of each functional run were dropped, and the frames that were flagged as motion-contaminated were removed and interpolated using a power-spectral matched interpolation (Power et al., 2014) and a temporal bandpass filter was applied from 0.009 to 0.08 Hz. Volumetric BOLD data was then mapped to each subject’s midthickness left- and right-hemisphere cortical surfaces generated by FreeSurfer from the atlas-registered T1 (Dale et al., 1999). The native surfaces were then aligned to the fsaverage surface in 32k fs_LR space and the two hemispheres were registered to each other using a landmark-based algorithm (Anticevic et al., 2012), such that they would be in geographic correspondence with each other to allow for point-to-point comparisons between each individual and across hemispheres (Glasser et al., 2013). Lastly, surface resting-state timeseries were spatially smoothed with a geodesic smoothing kernel (σ = 2.55, FWHM = 6mm). Functional connectivity was then calculated via temporal correlation between each vertex (a point on the cortical surface) timeseries with every other vertex timeseries.

### 2.4 Mapping locations of individual differences in brain networks

Locations of individual differences in functional network organization were identified using the “network variant” method as in (Seitzman et al., 2019). This method was chosen due to its ability to reliably identify regions of strong deviation in individual subjects, as opposed to regions that may differ only slightly in functional connectivity patterns. In addition to the results of the network variant method being robust, the network assignments of regions identified as variants have previously been validated using task-evoked activations (i.e. network variants show task-evoked activation in concordance with their assigned networks, and not their spatial location). **Figure 1** shows a schematic representation of the full variant identification and functional network assignment procedure.

**Figure 1.**
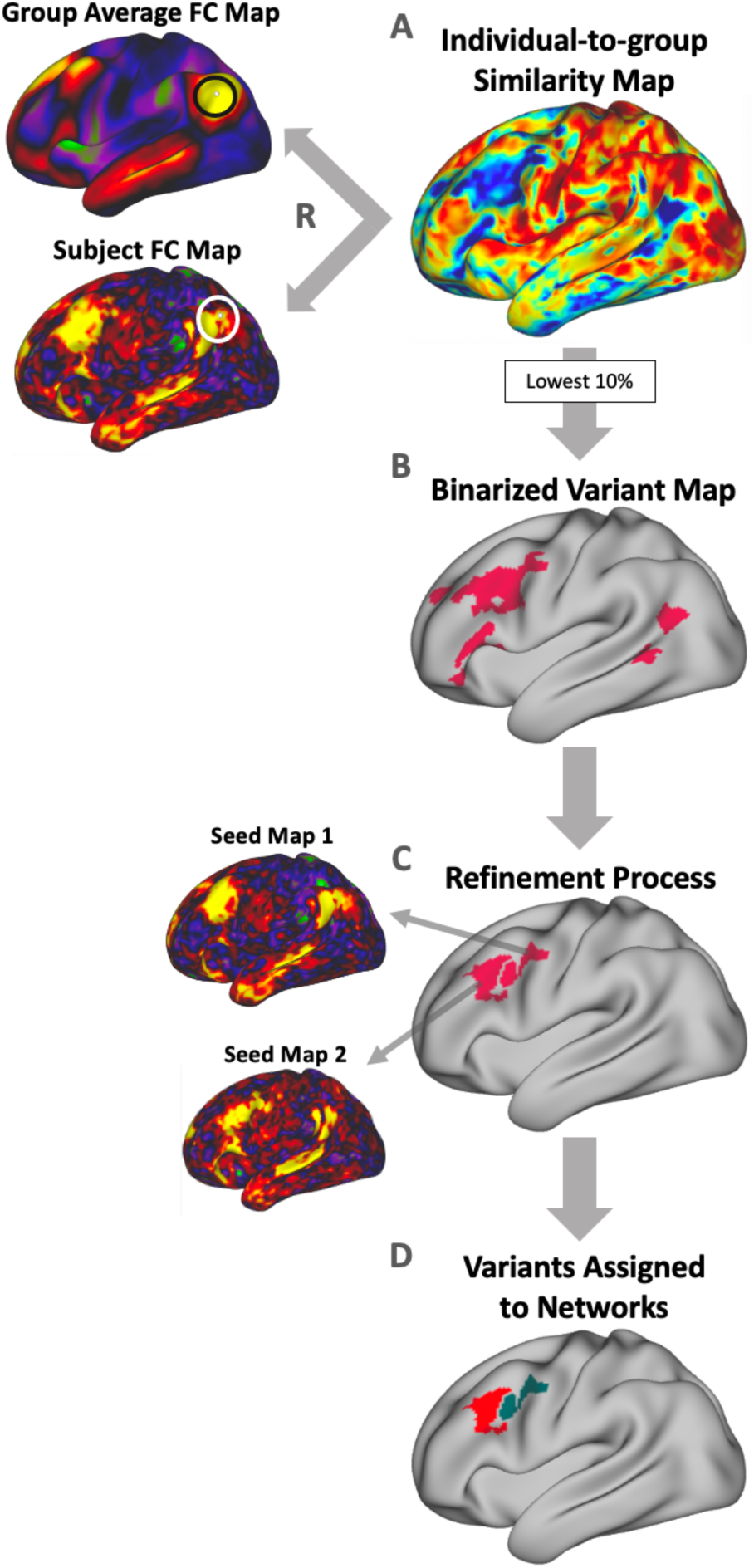
Procedure for identifying “network variants”, locations of strong individual differences in functional brain networks. Network variant regions are identified by comparing the functional connectivity of an individual to that of a group average derived from data from 120 young adults (WashU 120; see *Methods*). **(A)** First, a vertex-wise comparison of an individual-specific functional connectivity map and the group average (shown for a seed located near temporoparietal junction) is calculated using spatial correlation, resulting in an individual-to-group similarity map containing a continuous similarity value at each vertex. **(B)** The individual-to-group similarity map is then binarized to the lowest 10% of correlations to identify regions where the individual is most different from the group; small regions or regions with low BOLD signal are excluded. **(C)** These initial contiguous regions are then split into separable homogenous clusters as in (Dworetsky, Seitzman, Adeyemo, Smith, et al., 2021) (see *Methods*) and small clusters (<30 vertices) removed. Shown are average seed maps for two separable clusters that were split during the refinement process. **(D)** Lastly, each resulting variant region is assigned to the canonical network to which its average seed map is most similar, judged via Dice coefficient.

To begin, a cortical location-to-location (vertex-to-vertex) functional connectivity map was created for each subject based on their BOLD time series, concatenated across runs. Individual-to-group similarity maps were then obtained for each subject by comparing a given individual’s connectivity pattern at each vertex to the group average (WashU 120) connectivity pattern at that vertex using spatial correlation. Each subject’s similarity map was then thresholded to the lowest decile to identify the 10% of vertices that were most different between the individual and the group average. These vertices with the lowest correlation values were then binarized and small clusters (<50 vertices) were removed. Susceptibility areas (primarily along the inferior temporal cortex) with mean BOLD signal under 750 in the group average were set to 0 and excluded from analyses for all individual-specific connectivity maps due to poor signal quality.

In order to account for nearby, but distinct, network variant regions, these initial network variants were further refined into separable units as in (Dworetsky, Seitzman, Adeyemo, Smith, et al., 2021). This procedure takes into account the homogeneity of functional connectivity within a contiguous region, as well as the proportion of the variants’ territory that was dominated by a single network. Homogeneity was assessed using principal component analysis of a variant’s vertices’ seed maps, then calculating the variance explained by the first principal component of the variant (Dworetsky, Seitzman, Adeyemo, Smith, et al., 2021; Gordon et al., 2016). Network dominance was assessed via a vertex-wise template matching technique that assigned each vertex to the canonical network that was most similar to the vertex’s seed map. This similarity was calculated based on the Dice coefficient between the vertex’s seed map and each canonical network template, each binarized to the highest 5% of correlations (see (Dworetsky, Seitzman, Adeyemo, Smith, et al., 2021)). Variants that did not meet the threshold of 66.7% homogeneity and 75% network dominance in the individual network map were split along the network boundaries of a vertex-wise network map, producing the final analyzed network variants. These parameters for homogeneity and network dominance were set based on a pilot dataset (the Midnight Scan Club) for which manual ratings were used to flag variant clusters that appeared to be composed of multiple heterogenous subunits. The parameters used in the refinement process for the larger HCP subsample were the values that were better able to identify such clusters in the pilot dataset. Any resulting clusters after the refinement process that were smaller than 30 contiguous vertices were removed due to their small size. Previous analyses have shown that these parameters can identify large clusters that may consist of separate but adjacent network variants (Dworetsky, Seitzman, Adeyemo, Smith, et al., 2021). In addition, some supplemental analyses were conducted with pre-variants prior to this variant refinement procedure (**Appendix A**).

By definition, network variants are associated with different network patterns than what would normally be found at that location. In order to assess which large-scale network a variant was associated with, the resulting variants were matched to the best-fitting functional network using network templates from 14 canonical networks: default mode network (DMN), visual network, frontoparietal network (FPN), dorsal attention network (DAN), language network, salience network, cinguloopercular network (CON), somatomotor dorsal (SMd) and lateral (SMl) networks, auditory network, temporal pole network (T-pole), medial temporal lobe network (MTL), parietal memory network (PMN), and parietal occipital network (PON). Templates (or average connectivity maps) for each network were generated using data from the WashU 120 dataset and binarized by thresholding the top 5% of correlations (Gordon, Laumann, Gilmore, et al., 2017; Seitzman et al., 2019). For each network variant, the average seed map for all vertices within the variant was computed and also binarized to the top 5% of correlations. This binarized variant seed map was then compared in similarity to each of the binarized network template connectivity patterns via Dice coefficient (as in Dworetsky et al., 2021). The variant was then assigned to the network to which it was most similar. Network variants were removed if they did not match to any functional network (Dice ≤ 0) or if more than 50% of their territory overlapped with the territory of its assigned network in the group average. Note that, while here we use the label “language network” to refer to the network located on the superior temporal gyrus and a portion of the inferior frontal gyrus, this network has also been referred to as the “ventral attention network” (VAN) in previous reports. However, recent reports suggest that this network is more accurately described as a language network due to its correspondence with the expected distribution of language regions (Braga et al., 2020; Lipkin et al., 2022).

### 2.5 General Size and Number of Network Variants Across Hemispheres

The left and right hemispheres exhibit marked asymmetries at every scale of organization, thus, we asked if the properties of individual differences observed in functional connectivity patterns also vary across the two hemispheres. We first examined whether general properties of size and number of network variants differ across the two hemispheres. To this end, we compared the total number of network variants and average variant size between the left and right hemisphere. Network variant size was calculated by converting vertices belonging to each network variant to surface area using the wb_command function -surface-vertex-areas on the Conte69 group average midthickness cortical surface (Glasser & Van Essen, 2011). These were then compared between the left and right hemispheres. We also calculated a “magnitude” of asymmetry defined as the difference in the measurements of the two hemispheres divided by the greater value. This resulted in a percentage that indicates how much greater one hemisphere was compared to the other.

Because the values that we measure in this study are likely to exhibit a non-normal distribution, we used a non-parametric approach to test the significance of the observed differences. Permutation testing was used to test our hypothesis that the properties of variants differ across the two hemispheres by comparing the true difference values to a null distribution created by randomizing the hemisphere labels. Under the null hypothesis the hemisphere labels would be exchangeable because they do not differ significantly. Thus, in order to obtain a null distribution of difference values, , left and right hemisphere labels were permuted by subject by randomly selecting 50% of the subjects from the sample and switching the left and right hemisphere labels across 1000 permutations. For each permutation, we obtained the difference in the number of variants between the pseudo-left and pseudo-right hemispheres and averaged the differences across subjects’ permuted hemispheres. We repeated this procedure 1000 times in order to create the null distribution against which the true difference score was compared. The significance threshold was set at p < 0.025 at either end of the distribution for two-tailed tests. This same test was conducted to test the significance of the difference in average network variant size and overall variant territory. We used false discovery rate (FDR) correction for multiple comparisons across three tests: number of variant regions, overall variant territory, and average variant size.

### 2.6 Spatial Distribution of Variants Across Hemispheres

Since some large-scale networks are associated with relatively lateralized functions, we hypothesized that network variants might exhibit different spatial distributions across the hemispheres. We first conducted an omnibus test of the similarity in variant distributions between the left and right hemisphere. Variant maps were overlapped across subjects to quantify the frequency of variants at each vertex. The resulting overlap maps contained a value at each vertex for the proportion of participants who have a variant at that location. The variant frequency maps of the left and right hemispheres were then compared to each other by aligning each homotopic pair and then using spatial correlation to assess the overall (true) similarity of their spatial distribution of variants. Given that the two hemispheres were registered to each other, we define the homotopic pair of a given vertex as the vertex with the same index in the contralateral hemisphere. The significance of the similarity across the two hemisphere was assessed using a permutation approach, where the true similarity was compared to a null distribution of correlation values obtained by randomly switching the hemisphere labels. Under the null hypothesis that the two hemispheres do not differ significantly, we would obtain equivalent similarity values from comparisons of overlap maps from shuffled hemisphere labels as from comparisons of the true overlap between the two hemispheres. Our hypothesis was that the true similarity value between the hemispheres would be lower than the permuted similarity values, suggesting that the hemispheres are significantly different. As in previous examples, in each of 1000 permutations the left and right hemispheres labels were exchanged for 50% of the subjects that were randomly selected for each permutation. Then, a variant overlap map was created by summing the number of variants present at each vertex across participants. Using this overlap map, a correlation value was again obtained to quantify the overall similarity in variant frequency between the pseudo-left and pseudo-right hemispheres. Because we hypothesized that the left and right hemisphere would be significantly different in variant distribution, we then calculated how many of the correlation values from the 1000 permutations fell below the true correlation value to obtain a significance score. The significant threshold was set at p ≤ 0.025 at either end of the distribution for a two-tailed test. This analysis was replicated in nine MSC subjects (**Appendix B**).

We conducted an additional post-hoc test to identify the locations that were most different across the two hemispheres. For this test, a difference map was created by subtracting the frequency of variants at each vertex in the right hemisphere from its homotopic pair in the left hemisphere. Significant locations in this difference map were identified through a permutation-based approach with cluster correction to correct for multiple comparisons. Specifically, across 1000 iterations, the left and right hemisphere labels were permuted by subject and used to create a variant overlap map. For each permutation, a vertex-by-vertex difference map between the pseudo-left and pseudo-right hemispheres was created. We then thresholded each map at 5% difference (∼19 subjects in the HCP subsample of 384 subjects) and calculated the size of any clusters of variants that exceeded this threshold, creating a distribution of cluster sizes. Clusters in the true difference map were then compared to this permuted distribution of cluster sizes; clusters larger than 95% of the clusters obtained through permutation were interpreted as significant. Additional cluster thresholds (3% and 10% difference) were tested and included in **Appendix C** to show the robustness of results to this selection choice.

### 2.7 Networks Asymmetries Across Hemispheres

We then examined if specific networks showed asymmetries in the amount of cortical territory that was labeled as variants, or idiosyncratic regions, across the two hemispheres. To do this, we first identified each subject’s variant vertices and separated them by network to which they were assigned. Then, we calculated the surface area across variants using the *wb_command* function *-surface-vertex-areas* on the Conte69 group average midthickness cortical surface (Glasser & Van Essen, 2011). This variant surface area was then divided by the total surface area of the corresponding hemisphere to account for slight differences in hemisphere size. We then calculated a difference score (by network) for each subject, where the variant territory assigned to a network in the right hemisphere was subtracted from the variant territory that was assigned to the network in the left hemisphere. The difference scores for all participants were then averaged to obtain a mean difference score for each network. To test for significance, we again used a permutation approach, where we randomly switched the left and right hemisphere labels on a subject-level and recalculated average difference scores for the surface area of variant territory assigned to each network. This was repeated 1000 times to generate full null distributions for each network. We compared the true difference scores to these null distributions to assess significance. The significance threshold was set at p ≤ 0.025 at either end of the distribution for a two-tailed test. We again used FDR correction for multiple comparisons across 14 comparisons for each network.

To contextualize asymmetries in network variants relative to the patterns seen in the group average, we examined the symmetry of the size each network in a group average using the WashU 120 reference group (WashU 120; see **Appendix D** for results of this comparisons in two additional group averages: Midnight Scan Club and subsample of 384 HCP subjects). To do this, we calculated the difference between the proportion of surface area that each network accounts for across the two hemispheres. We then compared these group-average network asymmetries with the asymmetries seen in variant surface area in each hemisphere (calculated as described above).

### 2.6 Analysis of network variant asymmetries across handedness groups

We next investigated the potential relationship between hemispheric asymmetries in individual differences and dominant handedness, which is itself lateralized in most individuals and has been linked to other relatively lateralized functions such as language production (Knecht et al., n.d.; Perlaki, 2013). We repeated the previous analyses for left- and right-handers separately. HCP subjects in a sample of 752 individuals were divided into handedness groups according to their Edinburgh Handedness Inventory laterality coefficient (LQ; Oldfield, 1971). Left-handers and right-handers were determined to be so if their LQ was between -100 and -41 and 41 and 100, respectively. For each handedness group, we repeated the process of obtaining hemispheric difference scores for total variant territory, average variant size and number of variants. The hemispheric differences (left – right hemisphere) for all members of each handedness group were then averaged and compared between left and right handers. We assessed significance of the handedness effect using permutation testing. In this case, we permuted the handedness group labels. Handedness labels for all participants were shuffled, retaining the size of each handedness group (i.e., 40 random participants were labeled left handers and 670 participants were labeled right handers). For each permutation, we compared the difference scores between the pseudo-left and pseudo-right handers. This process was then repeated 1000 times to obtain a distribution of comparisons between groups. The true difference score was then compared to that null distribution to assess whether left and right handers differed from each other in the different variant properties described here. Under the null hypothesis that left- and right-handers are not significantly different, shuffling the handedness group labels would yield difference scores approximately equivalent to the true difference score between left- and right-handers. If handedness was associated with altered variant asymmetries, our hypothesis was that the true difference score would fall in the tails of the null distribution.

This process was also used to compare left- and right-handers in the network assignment of variants. To compare left- and right-handers in the spatial distribution of variants, we again used permutation testing. For each of 1000 permutations, we randomly selected 40 individuals to be labeled as pseudo-left handers and 670 individuals to be labeled pseudo-right handers and obtained variant frequency maps as described above. The frequency maps for left- and right-handers were then compared using spatial correlation in each permutation, creating a null distribution of similarity values. The true correlation value between the variant frequency maps of left- and right-handers were then compared to this null distribution to determine significance. The significance threshold was set at p ≤ 0.025 at either end of the distribution for a two-tailed test.

## 3. Results

Lateralization of structure, function and behavior are thought to occur as a consequence of multiple factors, including evolutionary, developmental, experiential, and pathological variables (see Toga & Thompson, 2003 for review). Hemispheric asymmetries exist at all levels of brain organization, including large-scale brain systems. Here, we examined whether hemispheric asymmetries are present for network variants: focal regions of high dissimilarity between an individual’s functional network pattern and that of a group average. Asymmetries in these idiosyncratic network variants could provide insight into the nature of individual differences in brain organization and how they arise. Thus, here we compare the properties of network variants between the two hemispheres.

The publicly available Human Connectome Project dataset was used for primary analyses in this manuscript. This dataset contains relatively high amounts of resting-state data for a large sample (N = 384) of young adults. The Midnight Scan Club, a smaller but highly sampled dataset (N = 9 individuals with 10 sessions each), was used to replicate findings of asymmetries in spatial distribution. We examined four properties of network variants. (1) First, we assessed asymmetries in the average size, number of variant regions and the overall variant territory in each hemisphere. (2) Next, we compared the spatial distribution of variants across hemispheres. (3) Then, the amount of variant territory associated with different networks was also compared across hemispheres. (4) Lastly, we provide a deeper examination of the relationship between network variant asymmetries and handedness, a prominent behavioral asymmetry.

### 3.1 Average frequency and size of network variants differ significantly across hemispheres

To better understand the nature of hemispheric asymmetries in individual differences in brain organization, we examined whether variants tend to be bigger or appear more frequently in one hemisphere over the other. On average, network variants were significantly bigger on the right hemisphere compared to the left (p < 0.001 based on permutation testing; **Fig. 2A**; left hemisphere = 236.1 mm^2^, right hemisphere = 285.3 mm^2^). There was also significantly more variant surface area in the right hemisphere compared to the left (p < 0.001; left hemisphere = 2564.5 mm^2^, right hemisphere = 2978.9 mm^2^; **Fig. 2B**). We found a small but significant difference in the number of variants in each hemisphere, with the left hemisphere showing slightly more variants than the right (p = 0.002, **Fig. 2C**; left hemisphere = 11.16 variants, right hemisphere =10.64 variants; however, this result, unlike the others, was affected by choices in processing the variant sub-units, see **Appendix A**). These results indicate that the right hemisphere exhibits more variable functional architecture, with bigger variants and more overall variant surface area. While variants in the left hemisphere were slightly more numerous, this effect was weaker and more affected by processing choices.

**Figure 2.**
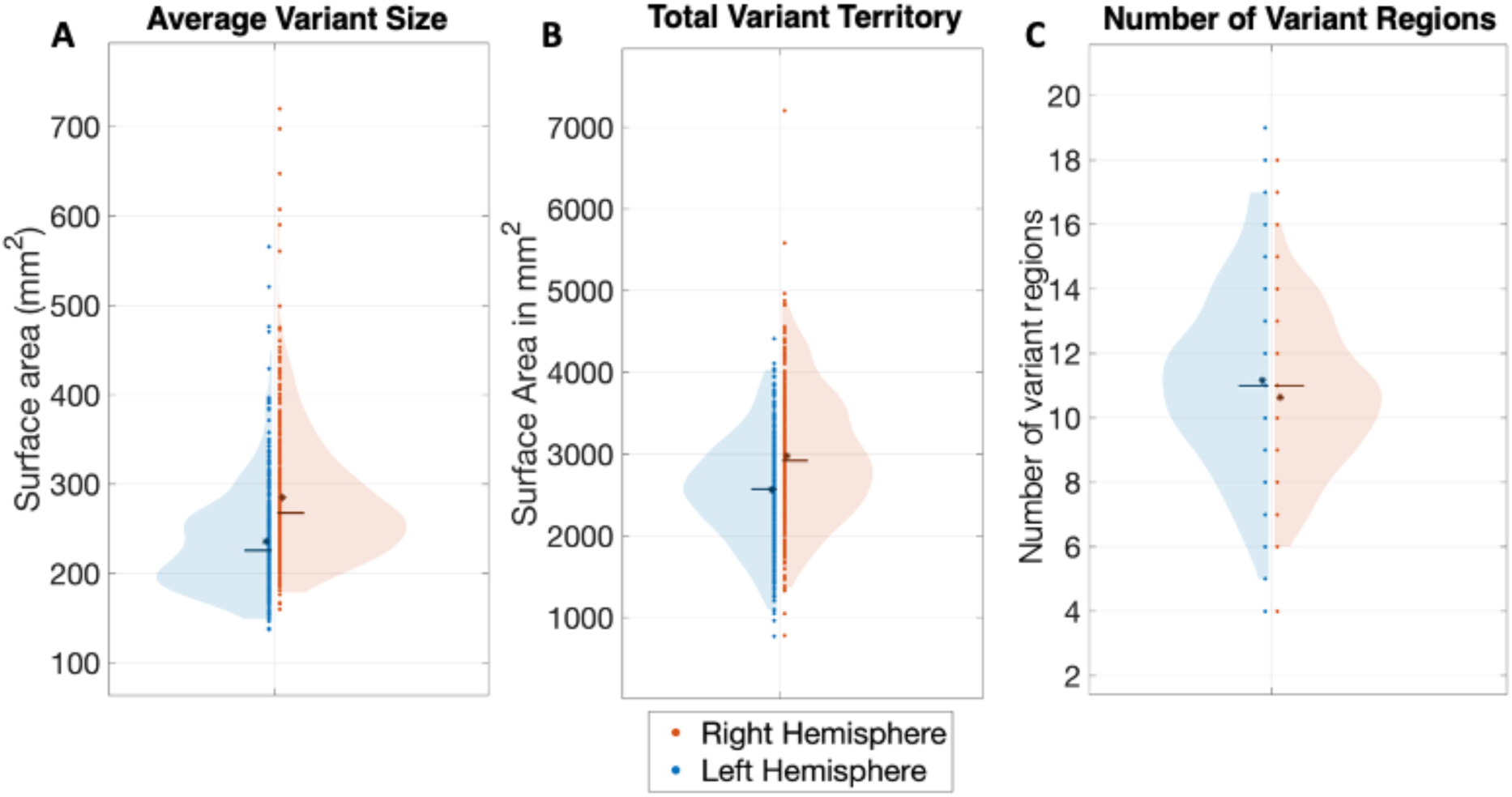
Comparison of the average number of separable variant regions and variant vertices, and average variant size across hemispheres. The violin plots show the distribution of measures of (**A**) average variant surface area (in mm^2^), (**B**) average total variant territory in the left hemisphere (left side of violin plot, blue) and the right hemisphere (right side of violin plot, red), and (**C**) average number of network variant regions. An asterisk indicates the mean and a line indicates the median of each distribution. Network variants in the right hemisphere were larger and covered significantly more surface area than in the left hemisphere. Left hemisphere variants were slightly more numerous (but this effect was somewhat more dependent on processing choices).

### 3.2 Spatial distribution of variants differs significantly across hemispheres

A relative lateralization can be observed in visualizations of the spatial distribution of network variants from past work (**Fig. 3A**; see also Fig. 3 in (Seitzman et al., 2019). This difference is most apparent in the lateral frontal cortex and near the temporo-parietal junction. Here, we tested whether this observed difference is significant by quantifying the similarity of variant spatial distributions between the left and right hemisphere.

Overlap maps of network variants were created for each hemisphere across subjects in the HCP dataset (**Fig. 3A**). The similarity in spatial distribution of network variants was compared between the two hemispheres using spatial correlation. This true similarity was then compared to those obtained across permuted comparisons (see *Methods*). The results of this analysis indicate that the spatial distribution of network variants differs significantly across the two hemispheres (**Fig. 3B**; p<0.001, true similarity between the left and right hemisphere: r = 0.74, mean permuted similarity: r = 0.99). This finding was replicated using the MSC dataset (**Appendix B**; p<0.001, true similarity: r = 0.39, mean permuted similarity: r = 0.52; note that here, correlations between all spatial distributions are substantially lower, likely due to the smaller sample size of the MSC dataset).

**Figure 3.**
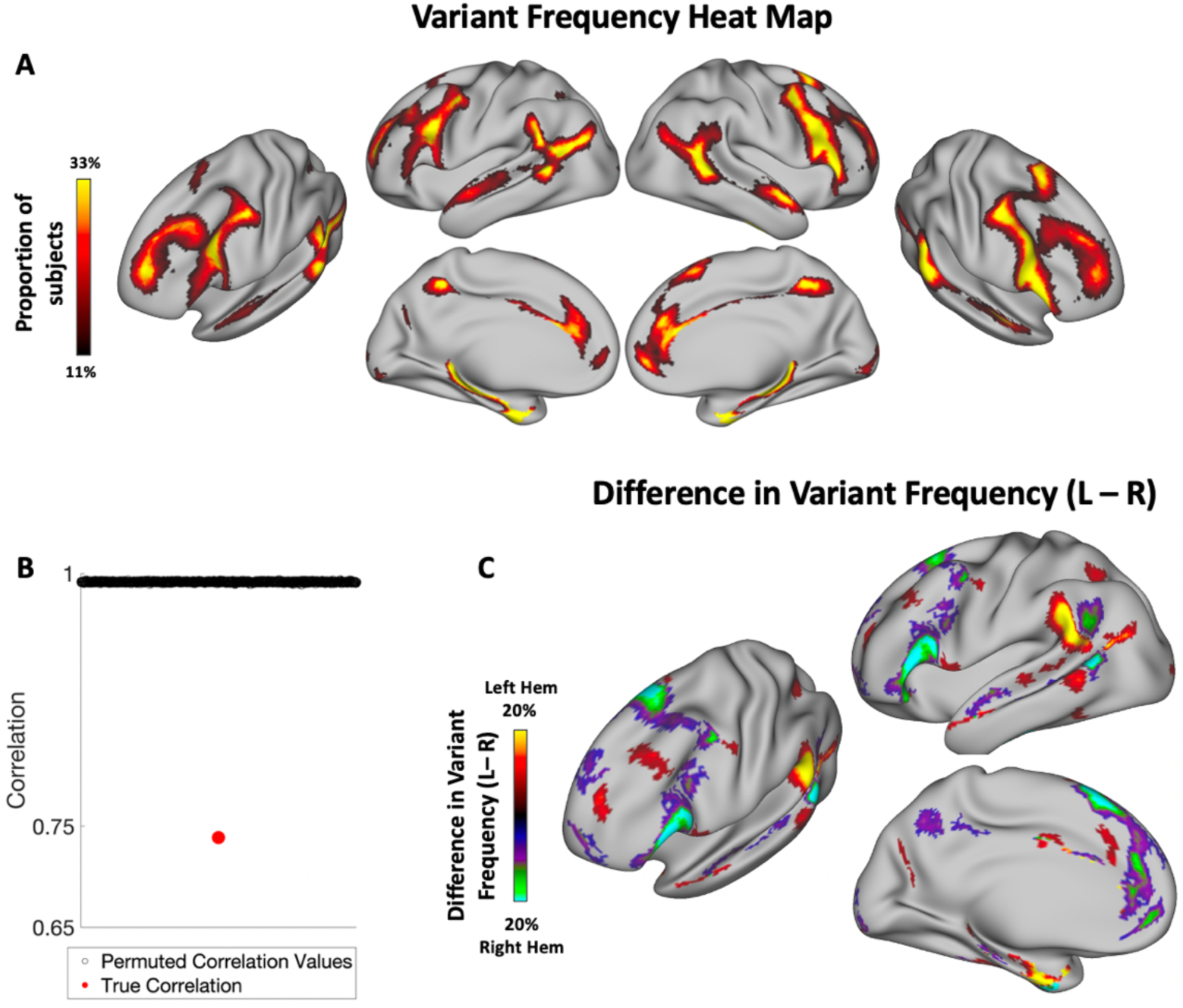
Hemispheric Asymmetries in the Spatial Distribution of Network Variants. **(A)** Frequency of network variants across 384 unrelated subjects from the Human Connectome Project. Network variants have a high frequency in regions of association cortex. **(B)** Permutation test examining the correlation between variant overlap maps of left and right hemisphere. Permuted correlation values indicate correlations between overlap maps obtained by randomly flipping the left and right hemispheres of subjects (see *Methods*); red dot indicates the true correlation. The two hemispheres have significantly lower similarity than expected by chance. **(C)** A difference map (projected on the left hemisphere for visualization) shows the regions in which the two hemispheres differ in the proportion of variant frequency, p<0.05 cluster-corrected at a frequency difference threshold of 5% (see **Appendix C** for other thresholds). Warm colors indicate a higher proportion of subjects that have a variant at that vertex in left hemisphere, while cool colors indicate a higher proportion of subjects that have a variant at that vertex in right hemisphere. The spatial distribution of network variants differs significantly across the two hemispheres, with the biggest differences observed in the inferior frontal gyrus, near the temporal parietal junction, and other localized areas of the frontal cortex.

To examine the specific locations where left and right hemispheres differ, we then did a second-level test where we subtracted the frequency of variants at each vertex in the right hemisphere from its homotopic vertex in the left hemisphere. A permutation-based cluster correction was conducted to identify locations of significant difference between the hemispheres (see *Methods*). This analysis shows that variants appear more frequently in the frontal operculum, angular gyrus, and superior medial frontal cortex in the right hemisphere and in the supramarginal gyrus, superior anterior frontal gyrus, and anterior portions of the medial temporal gyrus in the left hemisphere (**Fig. 3B**). This result was replicated using a smaller sample with high amounts of data per subject (MSC). The results of this replication analysis show a similar spatial distribution and regions of difference, though due to the small sample size, clusters were sparser (**Appendix B**).

### 3.3 Hemispheric asymmetries in the network assignment of variants

Network variants, by definition, are deviations from the canonical organization of a given functional network, associated with an atypical network for that location. Next, we explored how variants associated with specific networks differed between the hemispheres. First, each variant region was matched to one of 14 canonical networks using a template-matching procedure based on its seed map correlation pattern (**Fig. 1D**; see *Methods*). We then contrasted the extent of variants associated with each network in the left and right hemisphere.

Given the large asymmetries in variant territory between the two hemispheres, we focused first on whether variants of specific networks differed between the two hemispheres. Eight networks showed significant differences after FDR correction for multiple comparisons across 14 networks. Networks with more variant territory in the left hemisphere included the language network (p<0.004) and somatomotor lateral network (p<0.001), while those with more variant territory in the right hemisphere included the default mode (p<0.001), frontoparietal (p<0.001), dorsal attention (p<0.02), cingulo-opercular (p<0.001), parietal memory (p<0.004), and parietal occipital (p<0.001) networks (**Fig. 4**).Thus, hemispheric asymmetries exist in variants, but the direction of this asymmetry depends on the specific network. Namely, some networks are more likely to exhibit expanded idiosyncratic territory in the left hemisphere, such as the language and somatomotor network, while other higher cognitive networks are more likely to show idiosyncratic territory in the right hemisphere, like the default mode and cingulo-opercular networks.

**Figure 4.**
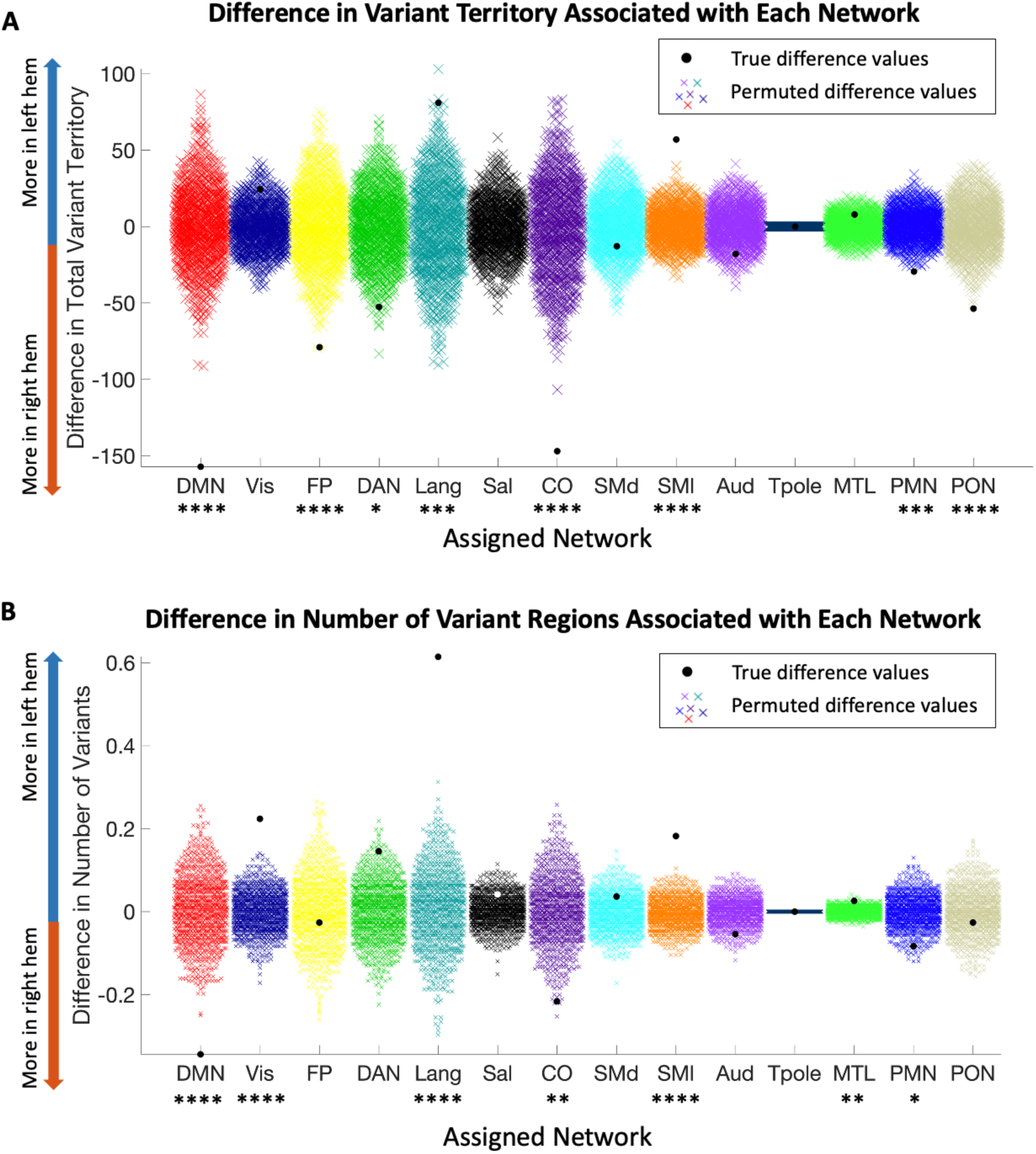
Hemispheric Differences in Network Assignment of Variants. A comparison of the A) total variant territory and B) number of variant regions associated with each network in the left and right hemispheres (color = network). Permutation testing shows that the language and somatomotor lateral have increased variant territory in the left hemisphere, while the default mode, frontoparietal, dorsal attention, cingulo-opercular, parietal memory, and parietal occipital networks have more variant territory in the right hemisphere. Similarly, the visual, language, somatomotor lateral, and medial temporal lobe networks have significantly more variant regions in the left hemisphere, while the default mode, cingulo-opercular, and parietal memory networks have significantly more variants in the right hemisphere than would be expected by chance. **** = p < 0.001; *** = p < 0.004, ** = p < 0.01; * = p < 0.02

Note that qualitatively similar results were also seen when asymmetries were compared for the number of variants, rather than territory covered. In this case, we found significant differences for 7 of networks. Variants were more commonly found in the left hemisphere for the visual (p<0.001), language (p<0.001), SMl (p<0.001) and MTL (p<0.01) networks, and in the right hemisphere for the DMN (p<0.001), CO (p<0.01), and PMN (p<0.02). Given the more robust findings associated with variant territory differences, we focus primarily on these comparisons moving forward.

Some of the networks that exhibit significant asymmetries in variants have been linked to functions or behaviors proposed to be differentially associated with the two hemispheres. For example, the language network has been shown to be left-hemisphere dominant in most individuals studied using precision approaches (Braga et al., 2020). Similarly, the DMN has been linked to episodic memory retrieval (Andrews-Hanna et al., 2010, 2014), a function that is hypothesized to be relatively right-lateralized (Desgranges et al., 1998; Tulving et al., 1994). Here, we looked in more detail at the spatial distributions of network variants for those that showed significant asymmetries (**Fig. 5**). Note that these maps are not cluster corrected for significance, but are instead used as an exploratory visualization of where the differences in variant for each network occur.

**Figure 5.**
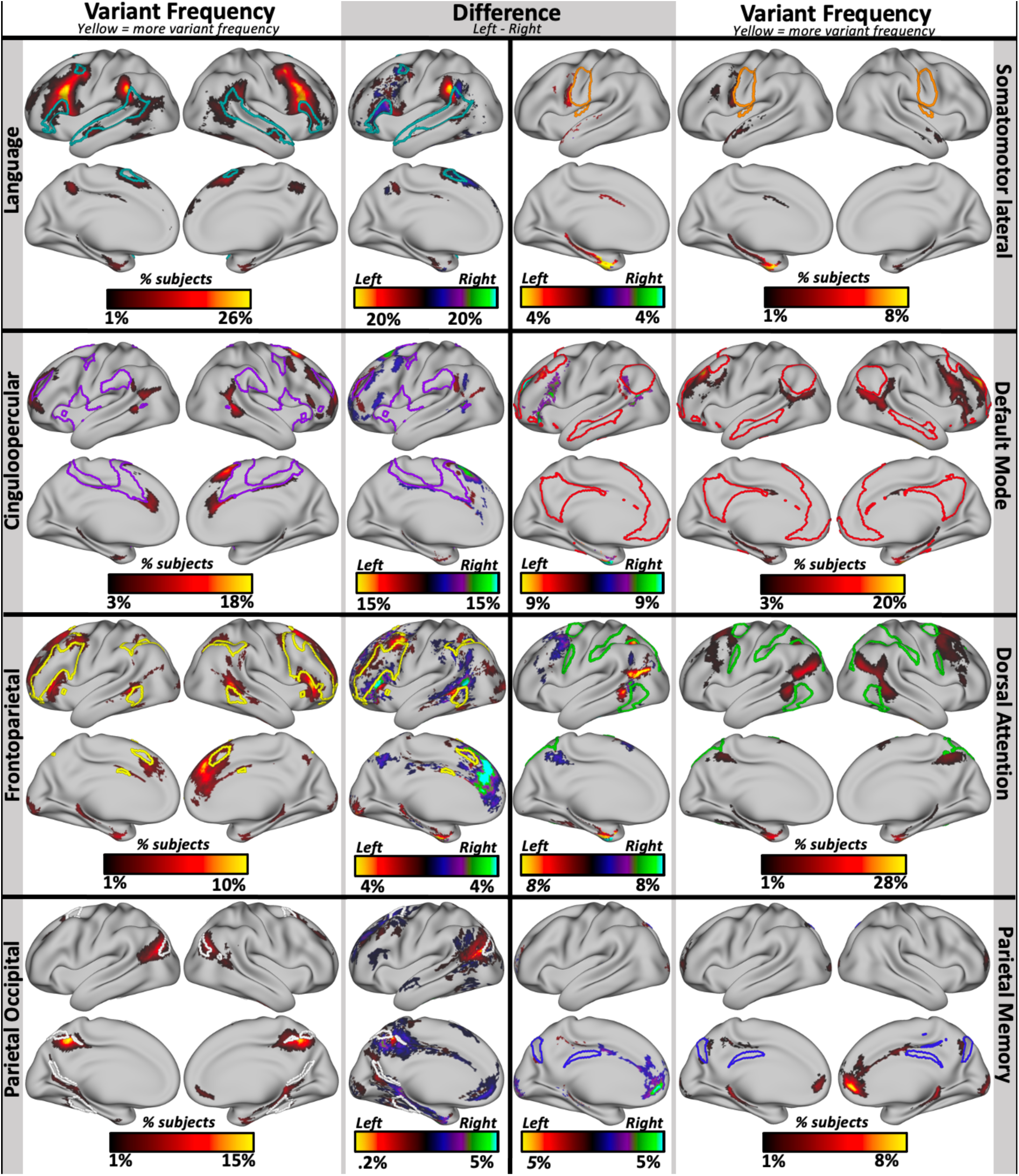
Areas of asymmetry in networks that showed significant hemispheric differences in their number of network variants. We examined the networks that showed significant hemispheric asymmetries in variant frequency in more depth to observe the regions where they differ the most across the two hemispheres. The language and somatomotor lateral networks have overall higher frequency of variant territory in the left hemisphere, while the cinguloopercular, default mode, frontoparietal, dorsal attention, parietal occipital, and parietal memory networks have higher frequency of variant territory in the right hemisphere. The lateral segments show variant frequency maps, where the color reflects the proportion of participants that have a variant for each network at that location. Note that the scale is different for each network to maximize visibility. The medial columns show difference maps where the right hemisphere frequency map was subtracted from the left hemisphere frequency map to show regions where the two hemispheres differ the most. Warm colors reflect higher variant frequency in the left hemisphere and cool colors indicate higher variant frequency in the right hemisphere. In each map, the colored outlines show the borders of the canonical network. Variant frequency and difference maps have not been cluster corrected and are presented primarily for qualitative comparisons.

The language network was associated with more variant territory in the left hemisphere especially on the supramarginal gyrus and pars opercularis, adjacent to group-average language network regions. The somatomotor-lateral network showed highest differences in a region on the inferior precentral gyrus, where variant frequency was significantly higher in the left hemisphere. The cinguloopercular network was associated with more variants in the right hemisphere, especially in a superior frontal area anterior to the precentral gyrus, further from its typical distribution. The default network had more variants in the right hemisphere, especially in regions adjacent to typical default locations along the superior frontal sulcus, and in some regions along the operculum and caudal pre-frontal cortex. The frontoparietal network shows a prominent difference in an anterior portion of the medial frontal gyrus, where it exhibits increased likelihood of variant territory in the right hemisphere compared to the left.

### 3.4 Relationship between asymmetries in network variants and network size

To contextualize the asymmetries in variants for each network, we asked whether there are asymmetries in the size of the networks found in the group average network map (**Fig. 6** **inset)**. We show that, even in the group-average, functional networks vary in the degree to which they show asymmetries (**Fig. 6A**). While most networks are relatively symmetric, the default mode and language networks were larger in the left hemisphere by 10%, equivalent to 1100 mm^2^, and 9%, equivalent to 349 mm^2^, respectively. The frontoparietal (yellow dot in **Fig. 6A**) and cingulo-opercular networks (purple dot in **Fig. 6A**) were larger in the right hemisphere by 14%, or 686 mm^2^, and 8%, or 606 mm^2^, respectively. Similar results were seen in group average networks from other datasets (**Appendix D**, e.g., note that across group averages the DMN consistently shows the same pattern of large difference, with more surface area in the left hemisphere).

**Figure 6.**
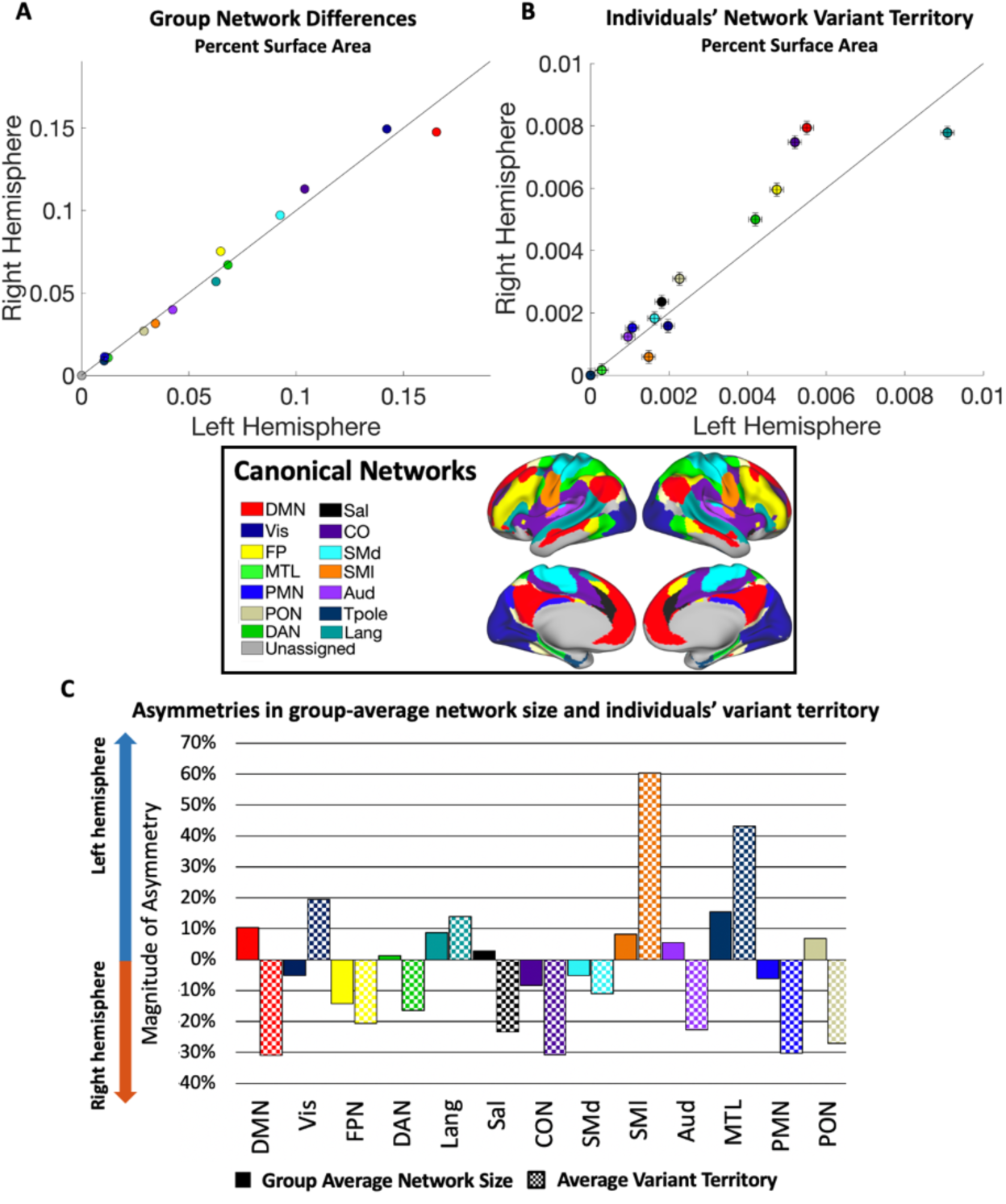
Asymmetries in group average network size and number of variants. **(A)** A comparison of group average network surface areas in left (x-axis) and right (y-axis) hemispheres (color = network, area expressed as a percent of total surface area). These group-average networks are largely symmetrical, with some small-scale differences. **(B)** Network variant territory associated with specific networks for the left (x-axis) and right (y-axis) hemispheres (error bars = SEM). **(C)** The magnitude of hemispheric asymmetries in size of group-average networks and variant surface area per network were quantified. The variant territory differs across the two hemispheres, sometimes in the same and sometimes in opposing directions from the asymmetries seen in the group-average maps.

These group-average network asymmetries can then be contrasted with the differences in variant surface area for each network (**Fig. 6B**). For example, if a network is relatively symmetric in the group average, but it exhibits more variant territory in one hemisphere compared to the other, this may suggest that individual differences in that network lead it to be more asymmetric within many people. This appears to be the case for the dorsal attention and parietal memory networks. Alternatively, if a network shows hemispheric differences in the group average, variants in one hemisphere could either magnify that asymmetry (if variants are ipsilateral to the group-average asymmetry), or it could counter it (if variants are instead more common in the contralateral hemisphere of the group-average asymmetry). We see apparent examples of both cases: for instance, cingulo-opercular and language variants occur primarily ipsilateral as their group-average asymmetries, whereas default mode variants appear primarily contralaterally (**Fig. 6C**). A similar pattern was observed when comparing the average number (rather than surface area) of variant regions associated with specific networks across the two hemispheres (**Appendix E**).

### 3.5 Relationship between handedness and hemispheric asymmetries

Handedness is a prominent behavioral trait that shows natural variance in the direction and degree of asymmetry throughout the population. Handedness has been shown to be related to other lateralized functions such as language. Because of this, we looked into asymmetries in network variants as a function of handedness in 40 left-handers and 670 right-handers from the HCP dataset (see *Methods* for handedness definition). The number, size, and spatial distribution of variants, as a whole, showed no significant differences between the two handedness groups. Interestingly, however, the two groups differed significantly in the specific networks that show asymmetries in their variants. Namely, we found significant interactions of handedness by hemisphere for two networks—the cinguloopercular (p = 0.004 based on permutation testing, FDR corrected for multiple comparisons) and frontoparietal network (p = 0.006, **Fig. 7A**)—two important networks for cognitive control. These interactions reveal that while right-handers do not differ significantly in their number of frontoparietal variants across hemispheres, left-handers show an increased number of frontoparietal variants in their left hemisphere (p = 0.006, **Fig. 7B**). In contrast, while right-handers have significantly more cinguloopercular variants in their right hemisphere (p < 0.001, **Fig. 7C**), left-handers do not show a significant difference. These findings add to the evidence suggesting that network variants are linked to behavior, and in particular, that asymmetries in network variants are related to asymmetrical functions and behaviors.

**Figure 7.**
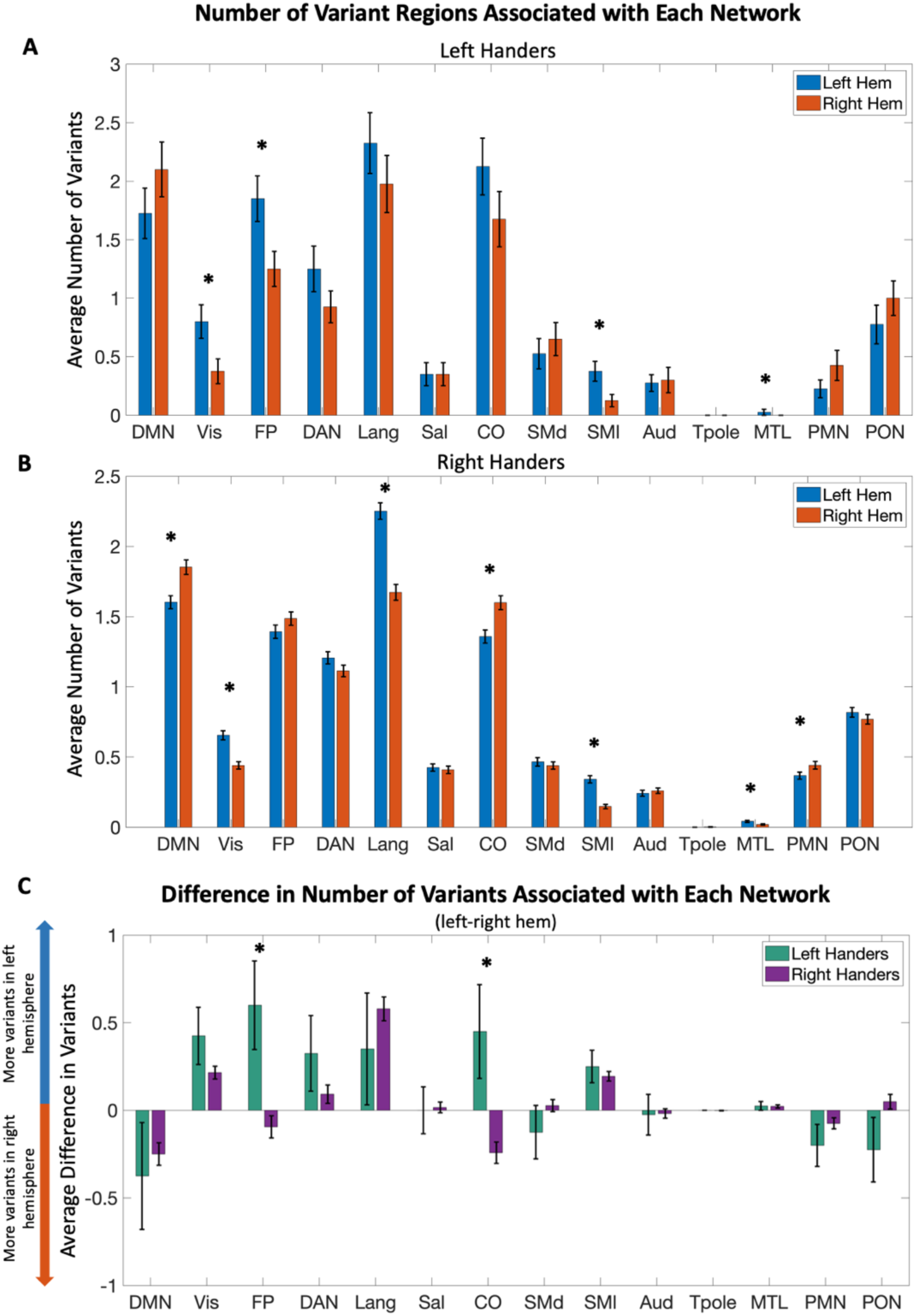
Difference in network assignment of variants across handedness groups. Comparison in the number of variants associated with each network across left and right hemispheres in **(A)** left-handers and **(B)** right-handers. Left-handed individuals showed significant differences in visual, frontoparietal, somatomotor lateral, and medial temporal lobe networks, with more variants in the left hemisphere. Right-handed subjects have significantly more right hemisphere variants associated with default mode, cinguloopercular and parietal memory networks, and more left hemisphere variants linked to visual, language, somatomotor lateral, and medial temporal lobe networks. **(C)** The bar graph shows the average difference across subjects in the number of variants associated with each functional network in their left minus their right hemisphere. Interactions between handedness and hemisphere were found for FP (p = 0.006) and CO (p = 0.002) network variants. These comparisons remained significant after FDR correction across all network comparisons. Error bars indicate SEM.

## 4. Discussion

The results described in this paper indicate that the properties of network variants differ across the two cerebral hemispheres. Generally, the right hemisphere shows larger variants, and more variant surface area. Variants were slightly more numerous in the left hemisphere, but this effect was small and dependent on specific pre-processing steps. These findings suggest that the right hemisphere exhibits a higher degree of variability in functional organization than the left.

A deeper examination of where specifically the two cerebral halves differ showed that the left hemisphere exhibits higher variant frequency on the supramarginal gyrus, superior anterior frontal gyrus, and anterior portions of the medial temporal gyrus, while the right hemisphere appears to have more variants on the orbital and triangular parts of the inferior frontal gyrus, angular gyrus, and superior caudal frontal cortex. These asymmetries also varied by network, with some networks exhibiting more variants in the left hemisphere, while others are more likely to have variants in the right hemisphere, and other networks showing a more even distribution of variants. While some of these asymmetries in network variants built on the asymmetries seen in group average network organization, others showed asymmetries in opposing directions. Finally, asymmetries in some networks (cinguloopercular, frontoparietal) varied between left and right handers, suggesting a link to functional traits. Jointly, these findings provide evidence that idiosyncratic network variants exhibit hemispheric constraints in their development. These constraints may be linked to differences in associated cognitive/behavioral functions.

### 4.1 Individual differences in brain networks add to or counter asymmetries seen in group averages

The distribution of network variants for eight functional networks differed across hemispheres, suggesting that these networks show increased variability in one hemisphere compared to the other^1^. Specifically, the default mode, cinguloopercular, frontoparietal, dorsal attention, parietal occipital, and parietal memory networks showed higher frequency of variants in the right hemisphere, and the language and somatomotor-lateral networks exhibited higher frequency of variants in the left hemisphere.

Our analyses in the group-average showed that cortical networks, on average, show slight differences in the amount of cortical territory that they cover across the two hemispheres. Thus, a hemispheric asymmetry in variants associated with a network may either magnify or counteract the typical ‘group-average’ pattern. For example, the language and cingulo-opercular networks both have more overall variant territory (and increased number of variants) in the ipsilateral hemisphere as their group-average asymmetry, suggesting that variants of these networks tend to manifest as a greater degree of lateralization in some individuals (this was also true to some degree for the somatomotor lateral network, though because this system is relatively small, the hemispheric differences in surface area in the group average were also small). In contrast, some networks exhibit more variant territory in the contralateral hemisphere relative to the group average dominance, such that the initial asymmetry in surface area may be regressed in some individuals. This appears to apply to the default mode and parietal occipital networks.

One network with expanded variants in the left-hemisphere was the language network. **Figure 6** shows that the language network appears to be slightly larger in the left hemisphere in group-averages. The relatively large number of language network variants in the left hemisphere points to two important aspects of this network: 1) it is consistent with prior observations that this network tends to be relatively left-lateralized in most individuals (Braga et al., 2020), and 2) that key regions of this network are highly variable across people (Fedorenko, 2012; Fedorenko & Blank, 2020). That group average representations do not strongly reflect the leftward lateralization of the language network indicates that the group average language network may be failing to capture important areas that exhibit a higher degree of variability across individuals (Dworetsky, Seitzman, Adeyemo, Neta, et al., 2021; Lipkin et al., 2022). These highly variable regions that are not reflected in the group average would then be labeled network variants, explaining the asymmetry of language variant territory found in our analyses.

In contrast, core regions of the default mode network exhibit high concordance across individuals (Dworetsky, Seitzman, Adeyemo, Neta, et al., 2021). This network also shows a relationship with putatively asymmetrical functions, such as episodic memory and retrieval (Desgranges et al., 1998; Tulving et al., 1994). As we show here, the default mode network is associated with increased surface area in the left relative to the right hemisphere in group averages. However, within single individuals this network showed expanded variant territory (and variant numbers) in the right hemisphere compared to the left. In contrast to the lateralization of the language network, the higher number of DMN variants in the right hemisphere may indicate a relative “re-normalization” of this network in some individuals. This network has been associated with a range of introspective functions including episodic retrieval, future planning, and social tasks like mentalizing, likely fractionating into separable sub-components (Andrews-Hanna et al., 2014; Buckner & DiNicola, 2019; DiNicola et al., 2020; Mars et al., 2012; Saxe & Kanwisher, 2003). The region where we observed a higher frequency of variants on the right hemisphere, around the angular gyrus, in particular has previously been implicated in theory of mind (Andrews-Hanna et al., 2014). Further research is necessary to confirm the implications of variant asymmetries for these different functions associated with the default mode network.

The frontoparietal network was associated with increased variant territory in the right hemisphere as well as an asymmetry of variants associated with this network in left-handers (though asymmetry was observed in the number of variants in the overall group). The frontoparietal network is an important cognitive control network that is highly integrated with other large-scale systems and is thought to provide rapid and flexible modulation of other networks (Marek & Dosenbach, 2018). This network has previously reported to show strong specialization within each hemisphere (Wang et al., 2014), and tends to sub-divide into left and right subnetworks in group maps (Smith et al., 2009), with the left hemisphere network showing early responses and the right-hemisphere network showing late onsets with prolonged responses during decision making (Gratton et al., 2017). Each sub-network also flexibly couples with either the default mode or dorsal attention networks (Spreng et al., 2010, 2012). Thus, it has been hypothesized that the left and right frontoparietal sub-networks contribute to different sets of functions through their interactions with distinct networks. The frontoparietal network is also linked to a particularly high number of variants relative to sensorimotor systems (Seitzman et al., 2019). An increased extent of variant territory associated with this network may suggest a larger link to rightward-asymmetrical functions. However, the finding that left-handers, unlike right-handers, show a significant asymmetry in this network, suggests that the frontoparietal network exhibits systematic differences across handedness groups that may be averaged out due to left-handers being a minority (and often excluded) from most samples.

### 4.2 Variant asymmetries inform us of developmental constraints on individual brain networks

What factors may give rise to asymmetries in idiosyncratic network variants? The cortical organization of the brain exhibits variation across individuals in the size, shape and spatial topography of functional networks. While size and spatial topography of networks has been shown to be moderately heritable (Anderson et al., 2021), variability in cortical regions may also arise due to developmental events that affect the way that functional systems are organized on the cortex. For example, signaling cascades direct the graded expression of transcription factors that regulate patterning of the cortex (O’Leary et al., 2007; Sur & Rubenstein, 2005) and alterations in their pattern of expression can result in alterations to the size and position of cortical areas (Fukuchi-Shimogori & Grove, 2001; Garel et al., 2003). Additionally, experiential factors have been shown to influence cortical organization. In cases of congenital blindness and deafness, the change in relative patterns of sensory-driven stimulation can lead to alterations in sensory domain allocation, cortical field size, and cortical and subcortical connectivity (Hunt et al., 2006; Kahn & Krubitzer, 2002; Striem-Amit et al., 2015). In congenital blindness, the absence of visual experience seems to lead to increase intra-individual variability in functional connectivity patterns, suggesting that sensory experience imposes some consistency in brain organization (Sen et al., 2021). In normative development, experience guides the development of face- and scene-responsive areas in the central and peripheral portions of the retinotopic map, respectively (Arcaro & Livingstone, 2017; Downing et al., 2001; Kanwisher et al., 1997; Srihasam et al., 2014). Interestingly, hemispheric dominance for certain functions is thought to arise as a result of competition for representational space. Due to proximity to language areas, the ventral occipito-temporal cortex in the left hemisphere is ideally situated for orthographic processing and as such becomes specialized for word perception, which may lead to a rightward asymmetry for face perception given lower competition in that hemisphere (Behrmann & Plaut, 2013; Dundas et al., 2015). Thus, the organization of brain regions and networks can be shaped by biological, environmental, and developmental factors.

A recent compelling hypothesis describes a potential mechanism that guides the development of functional networks at the individual level. The Expansion-Fractionation-Specialization hypothesis (DiNicola & Buckner, 2021) proposes that the disproportionate expansion of association areas in the human brain has provided large zones of cortex that share a distributed anatomical-connectivity motif, where frontal, temporal, and parietal areas are highly interconnected. Early in development, association cortex may exhibit a proto-organization characterized by a coarse network with poorly differentiated anatomical connectivity that, as developmental events occur and experience accumulates, fractionates into multiple networks that specialize in different functions (DiNicola & Buckner, 2021). Which function each network is associated with is determined through competitive activity-dependent processes but may be biased by differences in anatomical connectivity to regions that are relevant to its function (Braga & Buckner, 2017; Buckner & DiNicola, 2019). For example, language production regions may be “anchored” by orofacial motor regions important for speech due to their functional relationship and may therefore frequently develop adjacent to each other (Braga et al., 2020; Krubitzer, 2007). The result of this fractionation and specialization process are multiple fine-grained networks, with networks important for flexible cognitive functions being farthest from unimodal regions (Buckner & Krienen, 2013; Margulies et al., 2016).

Hemispheric asymmetries may reflect another form of specialization within this process. Transmodal functional systems that support flexible cognitive functions require integration between areas that are far apart (Mesulam, 1998), but inter-hemispheric connections incur extra processing costs. A hypothesis for the evolutionary advantage of lateralization in the central nervous system proposes that this motif arose to facilitate performing tasks in parallel, as well as fast processing for important functions such as language, which requires rapid sequential processing, and visuospatial processing, which requires rapid identification of objects and their relations (Güntürkün, 2017). This is supported by studies showing that both animals (Güntürkün et al., 2000; Rogers et al., 2004) and humans (Chiarello et al., 2009; Everts et al., 2009) perform better at doing tasks in parallel if they show increased lateralization of relevant functions. Thus, lateralization of networks may arise in order to decrease redundancy of processing (Levy, 1977; Vallortigara, 2006). This may be especially important for functions that require fast and serial processing, such as language. Therefore, the advantage of lateralization likely depends on the network and the processes that it supports.

Lateralization of functional networks may be biased by qualitative differences in the architecture of the hemispheres, such as differences in cortical microcircuitry. For example, pyramidal cell dendrites in the right hemisphere form on average more long-range connections compared to the left-hemisphere (Hutsler & Galuske, 2003). Indeed, the wiring patterns of the left hemisphere seem to be more suitable for specific, core linguistic processing than those observed in the right hemisphere, which in general is more interconnected (see Box 1 in (Jung-Beeman, 2005)). If a function or system becomes lateralized throughout development, a unique combination of genetic influences, developmental events, and idiosyncratic experiences will likely give rise to differences in the spatial topology of the network as it fractionates and specializes in the dominant hemisphere, giving rise to network variants that are more prominent in that hemisphere. Interestingly, association areas that show disproportionate expansion during evolution, and from infancy to adulthood (Hill et al., 2010), overlap significantly with areas of asymmetry in the degree of within-hemispheric functional connectivity (hemispheric specialization) (Wang et al., 2014). Thus, asymmetries in network variants may reflect a form of specialization of these functional systems that may arise as a consequence of qualitative differences in the way that the two hemispheres process information.

### 4.3 Asymmetries in individual differences may be markers for healthy and pathological differences in brain function

Previous research suggests that variability of cortical regions (e.g. (Schwarzkopf, 2011; Verghese et al., 2014)) and spatial topography of networks (Bijsterbosch et al., 2018; R. Kong et al., 2019) may have implications for behavior and cognition. Here, we found that asymmetries in two networks important for cognitive control, the cinguloopercular and frontoparietal networks, differed between left- and right-handers. Interestingly, the cinguloopercular network has been implicated in motor control in recent reports where it showed altered functional connectivity in response to disuse of motor circuits related to subjects’ dominant hand (Newbold et al., 2021). Thus, understanding the properties of network variants may help elucidate brain-behavior relationships. An important reason for characterizing brain-behavior relationships in normative samples is to understand how alterations may lead to different forms of pathology. The trait-like characteristics of variants and relationship to behavioral measures suggests their potential utility as biomarkers for atypical brain function associated with altered cognitive function.

Our findings demonstrate the existence of asymmetries in the properties of network variants in a sample of healthy young adults. However, some pathological conditions exhibit atypical asymmetries in brain function that could potentially alter the pattern of asymmetries observed in our sample. For example, autism spectrum disorder has been associated with decreased lateralization of language (Escalante-Mead et al., 2003; Floris et al., 2021; Jouravlev et al., 2020) as well as handedness and cortical structure in previous reports (Lindell & Hudry, 2013). This atypical language lateralization in autism has been shown to be largely independent of other asymmetries, as other large-scale systems (namely the default mode network and ‘multiple demand’ or frontoparietal networks) were not found to exhibit atypical asymmetries (Nielsen et al., 2014). Similarly, schizophrenia has been linked to differences in asymmetries of language and handedness (Artiges et al., 2000; Crow et al., 1996), as well as reduced anatomical asymmetries (Sommer et al., 2001). Thus, one theory for the cause of the disorder is delayed cerebral lateralization (Crow et al., 1996). This suggests that alterations in brain asymmetries may contribute to pathological conditions that may interfere with cognitive function. While the properties of network variants in clinical populations remain to be uncovered, investigating whether the pattern of asymmetry associated with individual differences described here would be altered in cases of atypical lateralization associated with the aforementioned conditions could potentially lead to identification of characteristics by which clinical populations can be stratified according to neurobiological profiles.

Hemispheric asymmetries may be altered by non-pathological factors as well, such as normative aging (Bäckman et al., 1997; Cabeza, 2002; Cabeza et al., 1997; Szaflarski et al., 2006). This decrease in hemispheric asymmetries may arise as a compensatory mechanism (Cabeza et al., 1997) or dedifferentiation processes (Li & Lindenberger, 1999). In addition to studying network variants and their asymmetries in clinical populations, it would be interesting to track their asymmetries throughout the lifespan. Changes in the asymmetries of network variants would suggest that their presence is under the influence of activity-dependent processes. Furthermore, understanding age-related changes to the properties of network variants could yield a better understanding of their potential application for therapeutic approaches and their relationship with lateralized functions and disorders. If network variants are trait-like features of brain organization that reflect cognitive differences across individuals, measuring their asymmetries and how they change as a function of pathological conditions and age, may serve to track disease processes and age-related cognitive decline and dementias.

### 4.4 Limitations and future directions

This work led to an increased understanding of how individual differences in functional connectivity compare across the two hemispheres. However, we note some limitations and opportunities for future investigations. First, the analyses described in this report found asymmetries in the frequency of variants in perisylvian regions. These regions have previously been found to be highly anatomically variable across individuals, though asymmetries tend to be small in magnitude (Van Essen, Glasser, et al., 2012). While we did not examine the relationship between anatomical variability and location of network variants in this study, previous reports have shown that the locations of network variants do not correlate with measures of gross anatomical deformations. Seitzman et al., (2019) showed that regions labeled as network variants do not overlap well with large deformations that occurred during surface registration within the individual, suggesting that network variants don’t systematically arise due to anatomical variability (Seitzman et al., 2019). Additionally, it has been shown that individual-specific features of functional organization persist even after controlling for accuracy of anatomical registration (Gordon, Laumann, Adeyemo, et al., 2017). Future work will be needed to determine if finer scale anatomical features, or more specialized anatomical-functional relationships, relate to asymmetries in network variants.

Second, the networks used for these analyses are an estimation based on the functional connectivity at rest of a group average. Their correspondence with task-evoked activations has not been verified at the individual level in this work, though previous evidence indicates correspondence between resting-state functional networks and task activation maps (Andrews-Hanna et al., 2014; Braga et al., 2020; Gordon, Laumann, Gilmore, et al., 2017; Smith et al., 2009; Tavor et al., 2016). It is unclear how our method for matching variants with networks would perform for an individual who diverges extremely from the group average spatial pattern (e.g., an individual with a rightward asymmetry for language). Thus, the network labels assigned to variants here should be taken cautiously and verified functionally in the future.

Lastly, our analyses on the relationship between variant asymmetries and handedness uncovered hemispheric differences in the variants of two functional networks that are implicated in cognitive control, but the two handedness groups did not differ in other properties of variant regions. While we relaxed our inclusion criteria (by including sets of related subjects) in order to increase our number of left-handers, our left-handed sample was only of 40 individuals, and the disparity in size between the left- and right-handed samples was substantial. Thus, the possibility remains that handedness may be related to hemispheric differences in other network variant properties that are more subtle and may require a larger sample of left-handed individuals to uncover.

This work provides a reference point for looking at potential altered asymmetries in network variants across various conditions. Future directions might include examining the patterns of network variants in individuals suffering with disorders that exhibit altered asymmetries in the brain and older adults, where the asymmetries observed in this report may show an atypical pattern due to pathological factors, experience accumulation, or dedifferentiation of functional systems. Lastly, network variants have not been examined in the cerebellum, but asymmetries of within-hemispheric connectivity show a mirrored pattern relative to that seen in the cerebrum (Wang et al., 2014). Thus, it would be interesting to examine if this mirrored pattern holds for asymmetries seen in cerebellar network variants.

## Conclusion

We examined hemispheric asymmetries in the properties of network variants, or regions in which patterns of functional connectivity differ strongly between an individual and a group-average representation. We found that, in general, the right hemisphere has more “variant territory”, which is linked to larger variant regions. Significant asymmetries were also found in the spatial distribution of network variants, which were more prominent around the inferior frontal gyrus and the inferior parietal lobule. These asymmetries varied by network, with some networks showing asymmetries in the same direction as asymmetries in group-average network patterns and others in the opposite direction. Finally, we found significant differences between left- and right-handed subjects in the asymmetries observed for variants of the cinguloopercular and frontoparietal networks, suggesting a relationship between network variant asymmetries and differences in behavioral traits. Jointly, these findings demonstrate that variant regions in large-scale functional networks differ systematically across the two hemispheres, indicating that they may be constrained by developmental differences and/or processes that result in functional hemispheric asymmetries. Furthermore, these findings in a sample of healthy young adults may serve as a benchmark to which we can compare future studies investigating asymmetries in network variants in conditions that have been shown to be associated with altered functional lateralization in the brain, such as aging, schizophrenia, and autism spectrum disorder.

## Data Availability Statement

The data used for these analyses is publicly available and may be accessed at http://www.humanconnectomeproject.org/data/ and at https://openneuro.org/datasets/ds000224. The code used to analyze the data may be accessed at https://github.com/dianaperez25/PerezEtAl_HemAsymmetries.

## Acknowledgments

This research was supported in part through the computational resources and staff contributions provided for the Quest high performance computing facility at Northwestern University which is jointly supported by the Office of the Provost, the Office for Research, and Northwestern University Information Technology.

## Citation Diversity Statement

Recent work in several fields of science has identified a bias in citation practices such that papers from women and other minority scholars are under-cited relative to the number of such papers in the field (Dworkin et al., 2020; Fulvio et al., 2021). Here we sought to proactively consider choosing references that reflect the diversity of the field in thought, form of contribution, gender, race, ethnicity, and other factors. First, we obtained the predicted gender of the first and last author of each reference by using databases that store the probability of a first name being carried by a woman (Dworkin et al., 2020; Zhou et al., 2020). By this measure (and excluding self-citations to the first and last authors of our current paper), our references contain 10.46% woman(first)/woman(last), 11.89% man/woman, 22.62% woman/man, and 55.02% man/man. This method is limited in that a) names, pronouns, and social media profiles used to construct the databases may not, in every case, be indicative of gender identity and b) it cannot account for intersex, non-binary, or transgender people. Second, we obtained predicted racial/ethnic category of the first and last author of each reference by databases that store the probability of a first and last name being carried by an author of color (Ambekar et al., 2009; Sood & Laohaprapanon, 2018). By this measure (and excluding self-citations), our references contain 9.01% author of color (first)/author of color(last), 14.09% white author/author of color, 20.07% author of color/white author, and 56.83% white author/white author. This method is limited in that a) names and Florida Voter Data to make the predictions may not be indicative of racial/ethnic identity, and b) it cannot account for Indigenous and mixed-race authors, or those who may face differential biases due to the ambiguous racialization or ethnicization of their names. We look forward to future work that could help us to better understand how to support equitable practices in science.

## Appendix

**Appendix A.**
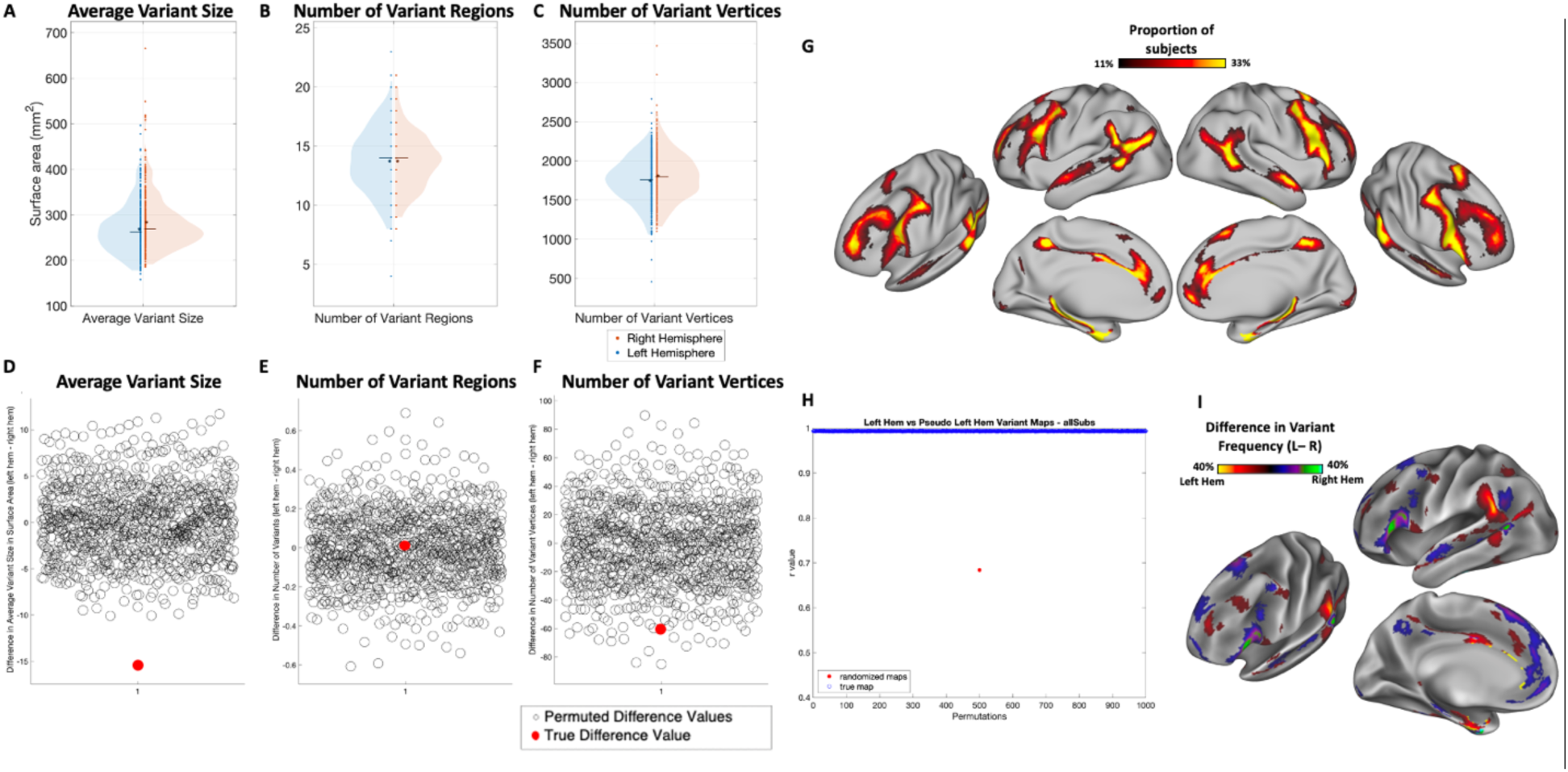
Examination of hemispheric asymmetries in the properties of network variants before undergoing the refinement process (“pre-variants”) A-C) Comparisons of average variant size, number of variant regions, and number of variant vertices across hemispheres in a subsample of 384 subjects from the Human Connectome Project. D-F) Results of permutation testing for significance. True difference value indicates the difference between *the true* left and right hemispheres. Permuted difference values indicate differences obtained by randomly flipping the left and right hemispheres of subjects 1000 times. G) Network variant overlap across subjects. H) Results of permutation testing for significance. True map indicates the correlation between variant overlap maps of *the true* left and right hemisphere*s*. Randomized maps values indicate correlations between overlap maps obtained by randomly flipping the left and right hemispheres of subjects 1000 times. I) A difference map shows the regions in which the two hemispheres differ in the proportion of variant frequency. Warm colors indicate more variant overlap in left hemisphere, while cool colors indicate more variant overlap in right hemisphere. The spatial frequency of network variants differs significantly across the two hemispheres. Although this analysis was conducted on network variants that were defined differently than those in the main text, the general distribution and differences across the hemispheres are replicated (compare this figure with Figure 3).

**Appendix B.**
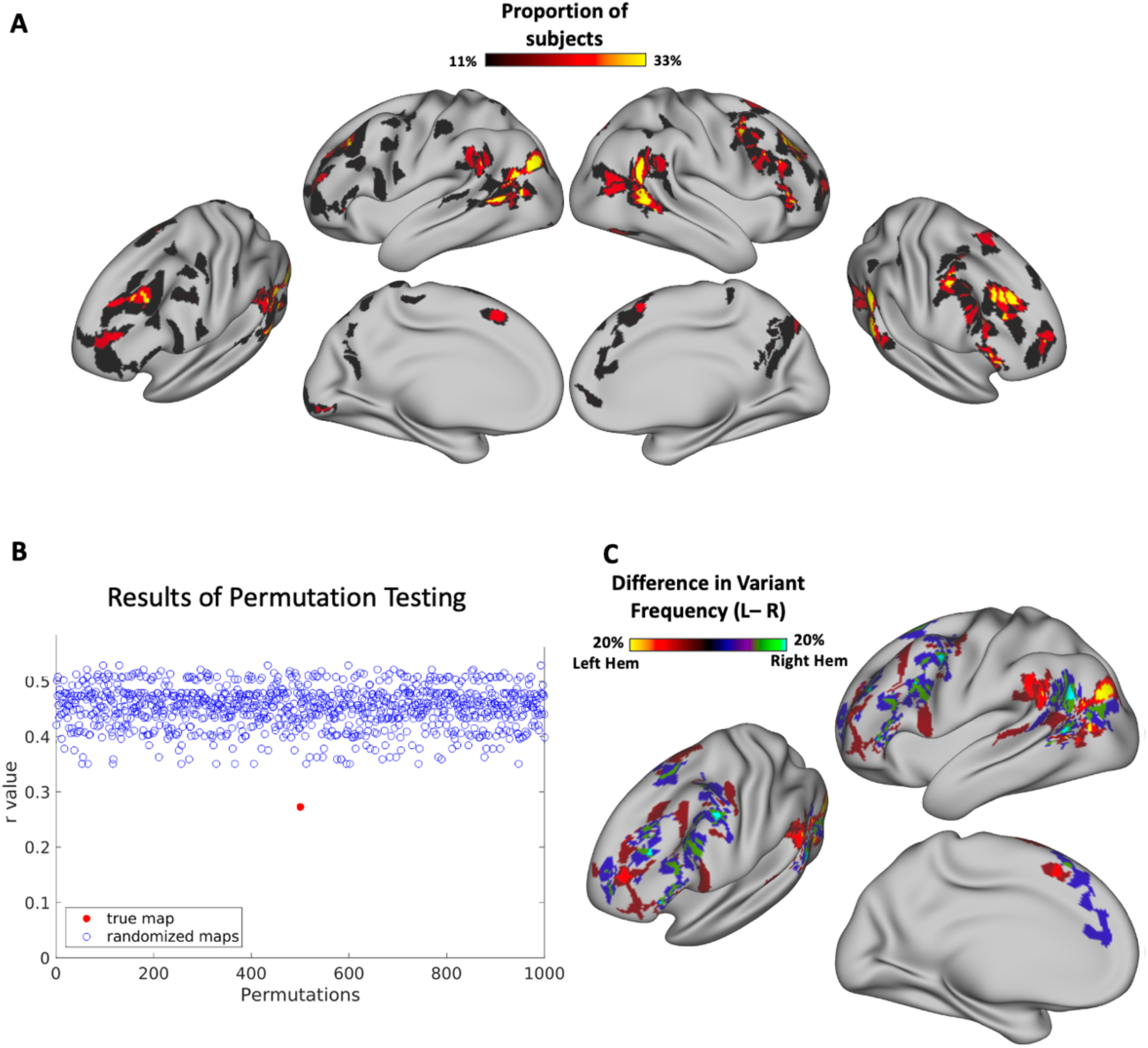
Examination of hemispheric asymmetries in the spatial distribution of network variants in an independent sample (Midnight Scan Club; MSC) A) Network variant overlap across 9 subjects from the Midnight Scan Club. B) Results of permutation testing for significance. True map indicates the correlation between variant overlap maps of *the true* left and right hemisphere*s*. Randomized maps values indicate correlations between overlap maps obtained by randomly flipping the left and right hemispheres of subjects 1000 times. C) A difference map shows the regions in which the two hemispheres differ in the proportion of variant frequency. Warm colors indicate more variant overlap in left hemisphere, while cool colors indicate more variant overlap in right hemisphere. The spatial frequency of network variants differs significantly across the two hemispheres. Although this analysis was conducted on a substantially smaller number of participants, with very different spatial and temporal scanning parameters, the general distribution and differences across the hemispheres are recapitulated (compare this figure with Figure 3). The substantially smaller sample size likely accounts for sparser clusters.

**Appendix C.**
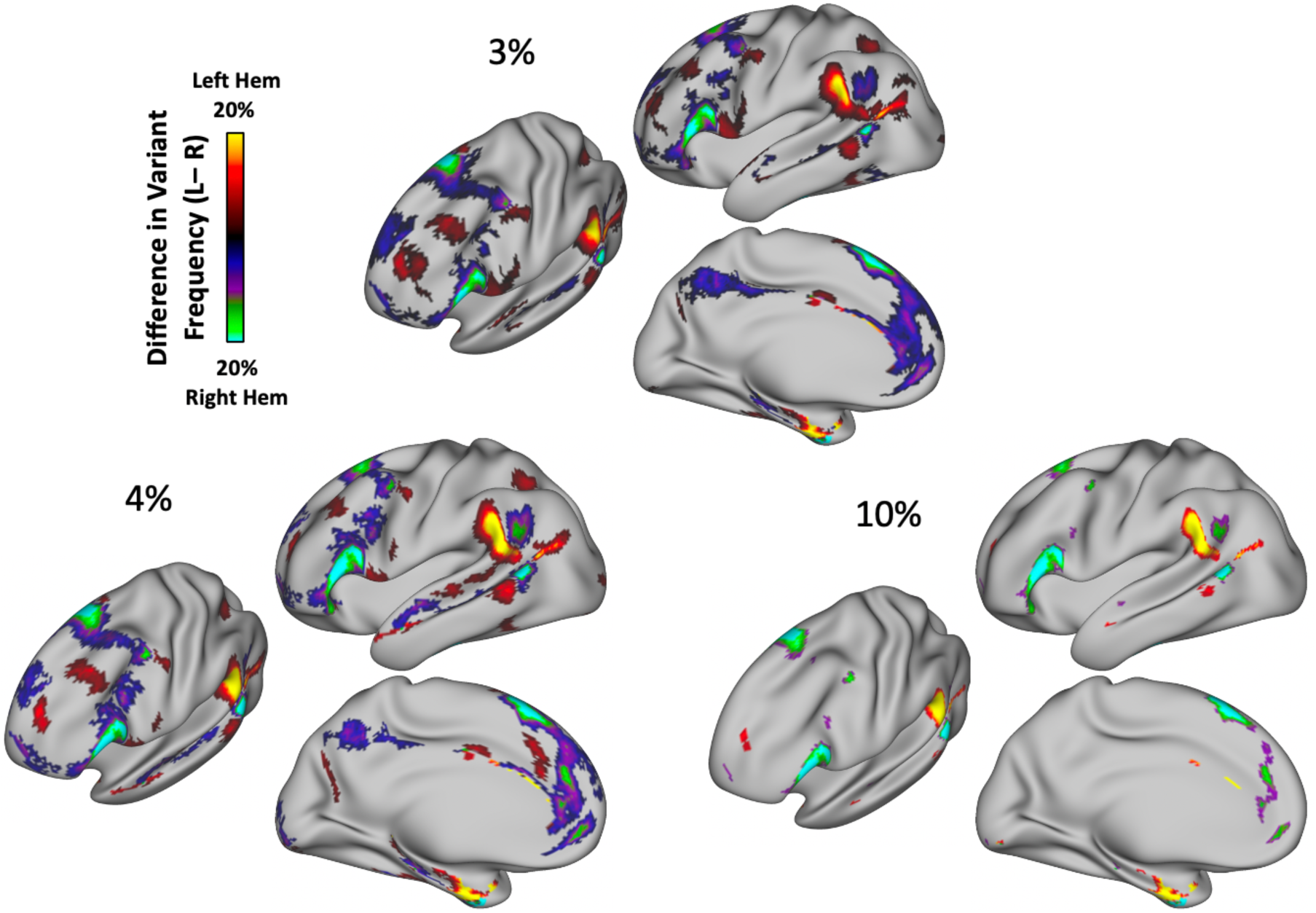
Comparison of cluster correction thresholds. Three cluster difference thresholds (3, 4, and 10%) were tested to observe the effect of this parameter on the results. The asymmetries seen across the two hemispheres appear to be robust to the effects of this parameter; smaller clusters are lost as cluster difference thresholds are raised. For the results described in the main text, a threshold of 5% was used.

**Appendix D.**
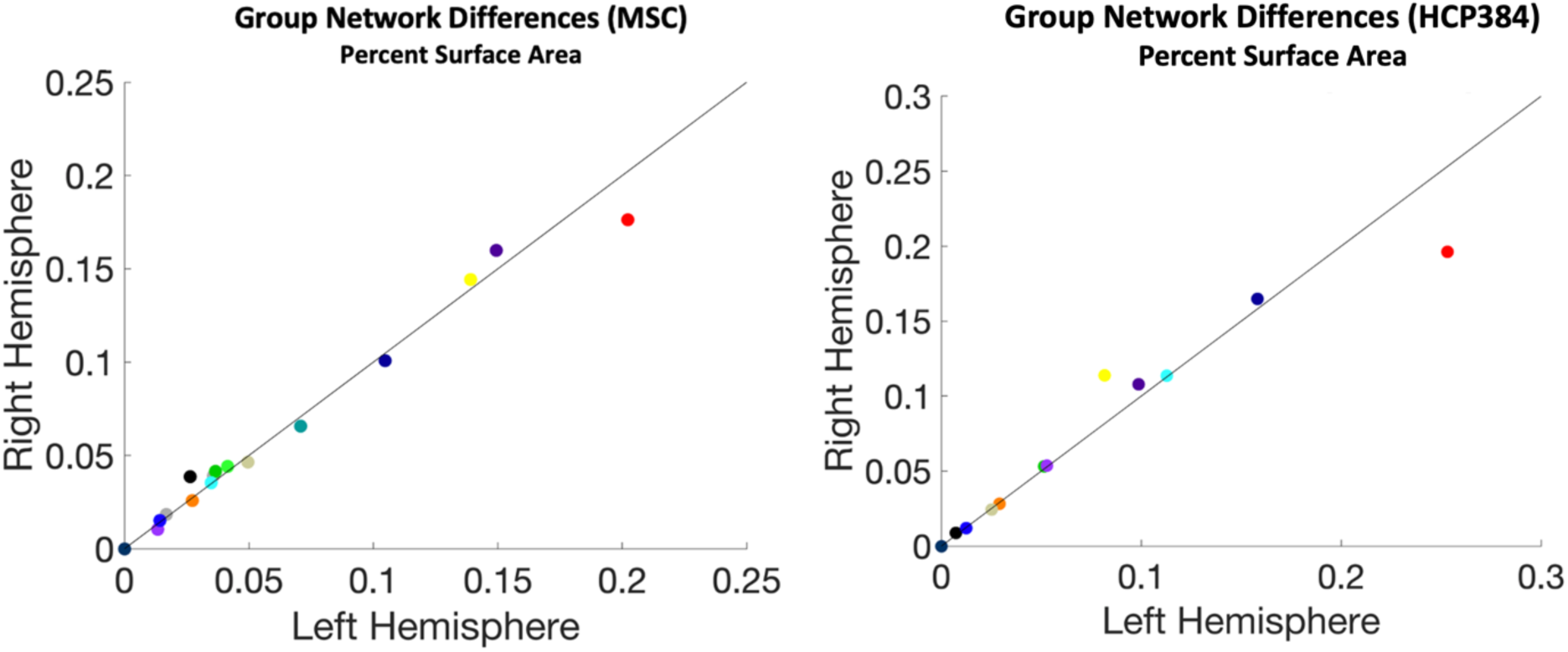
Comparison of surface area across hemispheres of *additional* group average networks. Two group averages—the Midnight Scan Club (MSC) and Human Connectome Project (HCP; n = 384)—were examined to observe the consistency of the pattern observed in the Wash U 120 group average (see Fig. 4). The default mode network consistently exhibits the biggest difference in surface area, with a left hemisphere lateralization (2.6% in MSC, 5.7% in HCP). Networks were defined using the data-drive network identification procedure InfoMap. Note that the HCP does not include the language, MTL or T-pole group-level networks because these networks do not emerge consistently across edge density in this dataset.

**Appendix E.**
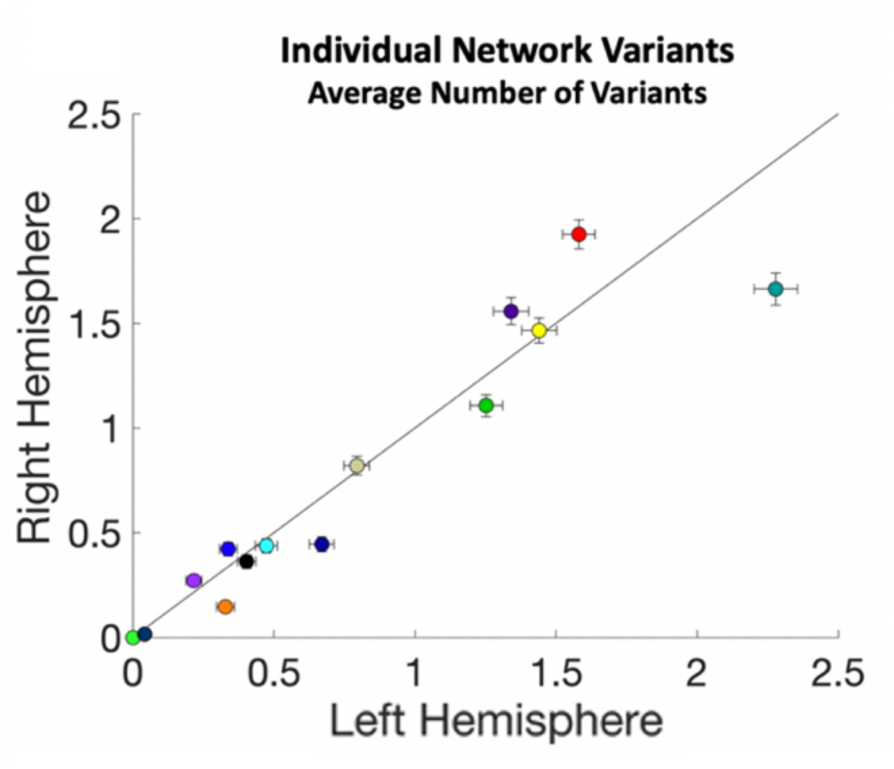
Comparison of number of network variant regions across hemispheres in the Human Connectome Project. The average number of variant regions associated with specific networks in the left (x-axis) and right (y-axis) hemispheres measured in the Human Connectome Project (HCP; n = 384). Each color corresponds to a different network. Error bars = SEM.

## Footnotes

1 Notably, our analyses did not test if the spatial location of regions was directly homologous; most analyses compared omnibus properties of the two hemispheres. In one case, we tested the spatial point-by-point correspondence of idiosyncratic variants between the two hemispheres; however, network variants do not necessarily correspond to distinct regions, and thus the presence of variant correspondence across the hemispheres cannot alone be taken as conclusive evidence of homology.

